# The rapid evolution of flagellar ion-selectivity in experimental populations of *E. coli*

**DOI:** 10.1101/2021.01.26.427765

**Authors:** Pietro Ridone, Tsubasa Ishida, Angela Lin, David T Humphreys, Eleni Giannoulatou, Yoshiyuki Sowa, Matthew A. B. Baker

## Abstract

Determining which cellular processes facilitate adaptation requires a tractable experimental model where an environmental cue can generate variants which rescue function. The Bacterial Flagellar Motor (BFM) is an excellent candidate – an ancient and highly conserved molecular complex for propulsion which navigates bacteria towards favourable environments. In most species, rotation is powered by H^+^ or Na^+^ ion transit through the torque-generating stator subunit of the motor complex. The ion that drives the rotor has changed over evolutionary timescales but the molecular basis of this selectivity remains unknown.

Here we used CRISPR engineering to replace the native *Escherichia coli* H^+^-powered stator with Na^+^-powered stator genes and report the rapid and spontaneous reversion of our edit in a low sodium environment. We followed the evolution of the stators during their reversion to H^+^-powered motility and used whole genome and transcriptome sequencing to identify both flagellar- and non-flagellar-associated genes involved in the cell’s adaptation. Our transplant of an unfit protein and the cells’ rapid response to this edit demonstrates the adaptability of the stator subunit and highlights the hierarchical modularity of the flagellar motor.

## INTRODUCTION

Bacterial motility via the flagellar motor represents one of the earliest forms of locomotion (Miyata et al., 2020). This rotary motility imparts such significant selective advantage (*1, 2*) that resources are allocated to chemotaxis even in the absence of nutrient gradients (*3, 4*). The evolutionary origins and subsequent adaptation of the motor are of significant scientific and public interest (*5*), since the BFM holds prominence as an ancient and large molecular complex of high sophistication. Furthermore, the BFM is an ideal model for studies in molecular evolution since it demonstrates modularity (*6, 7*) and single nucleotide variants which result in changes in motility are easily experimentally selected (*8*).

The torque that drives the BFM is supplied by motor-associated transmembrane protein-complexes known as stators. The stator complex, an asymmetric heteroheptamer (in *E. coli:* MotA_5_MotA_2_) most likely acts itself as a miniature rotating nanomachine coupling ion transit to rotation (*9, 10*). The stators are essential for motility, as they drive rotation, and are accessible for studies in experimental evolution due to their unambiguous role in connecting a specific environmental cue (presence of the coupling ion) to an easily discernible phenotype (cell swimming). Furthermore, the stators have been subject to protein engineering approaches for many years, in particular the synthesis of chimeric stator constructs that enable the motor of *E. coli*, natively proton-driven, to be powered by sodium ion flow (*11–14*). The majority of stators are proton driven, but many that are sodium-driven can be found in nature (*15*), and this divergence is presumed to have occurred in the distant past (*6, 16, 17*). Past reports have argued that H^+^-coupled motility diverged from Na^+^-coupled machinery in ancestral times (*18*) but the molecular basis for this adaptation, and the evolutionary landscape that constrains stator adaptation remains unclear.

In order to simulate the effects of natural evolution on stator adaptation we designed an experiment where an *E. coli* strain, expressing only a sodium-powered stator, would be introduced to a non-lethal environment (soft agar swim plate) which lacked the power source for the stator (Na^+^). Our hypothesis was that the population would undergo selection for upmotile variants, adapting its stators to function in the new environment.

We used genomic editing techniques (no-SCAR CRISPR/Cas9 (*19*) and λ-Red (*20*)) to replace the native *motA motB* stator genes of the *E. coli* BFM with chimeric sodium-powered *pomA potB* (henceforth Pots) stator genes derived from *Vibrio alginolyticus* (*11*). We transplanted the *pomApotB* stator genes at the same location and orientation of the native *motAmotB* locus to preserve the native genomic context of the motile RP437 *E. coli* strain. We then examined which genetic changes occurred during growth on soft-agar in depleted sodium, that is, under selective pressure for proton-driven motility. We performed our directed evolution experiments of our Pots *E. coli* strain in the absence of antibiotics to avoid additional, undesired selective pressures (*21*).

## RESULTS

### Directed evolution of Pots on low sodium swim plates

The RP437 strain was edited to carry the Pots stator genes in place of the native *E. coli motAmotB* genes via the no-SCAR Method (*19*) and traditional λ-Red recombineering respectively (Supplementary Fig. 1 & 2). A single no-SCAR Pots clone was selected and tested on swim plates (Supplementary Fig. 3) after verification of successful editing by colony PCR and Sanger Sequencing (Supplementary Fig. 4). The edited strain was able to swim on sodium-rich (Na^+^LB: ~100 mM NaCl) soft agar plates but not on potassium-rich sodium-poor swim plates (K^+^LB: 67 mM KCl, ~15 mM [Na^+^]) (Supplementary Fig. 3A). This edited strain exhibited the same swimming behaviour as the control stator-less strain with motility restored via an inducible plasmid vector that could express the Pots construct (RP6894 Δ*motAmotB* + pSHU1234, hereby pPots).

We next challenged this Pots strains to survive on K^+^ based soft agar (K^+^LB) for prolonged periods (Supplementary Fig. 3B). Motile subpopulations arose spontaneously from inoculated colonies within a few days. Cells from the edge of these motile flares were passaged onto fresh swim agar for up to 5 passages at 3-4 days intervals (Supplementary Fig. 5). When multiple flares occurred in a single swim ring, each was individually passaged (Supplementary Fig. 3B), and could be recapitulated (Fig. 1C, Supplementary Fig. 3C). Directed evolution consistently generated swimming flares when Pots clones were cultured on agar containing yeast extract and tryptone (K^+^LB swim plate: ~15 mM [Na^+^]), but not on minimal media (K^+^MM swim plate: ~1mM total [Na^+^]) or when the Pots construct was encoded on a plasmid (Supplementary Fig. 6). One Pots strain generated using λ-Red methods (*20*), which carried the native *V. alginolyticus* Shine–Dalgarno (SD) sequence, also successfully produced flares (Supplementary Fig. 7).

**Figure 1.**
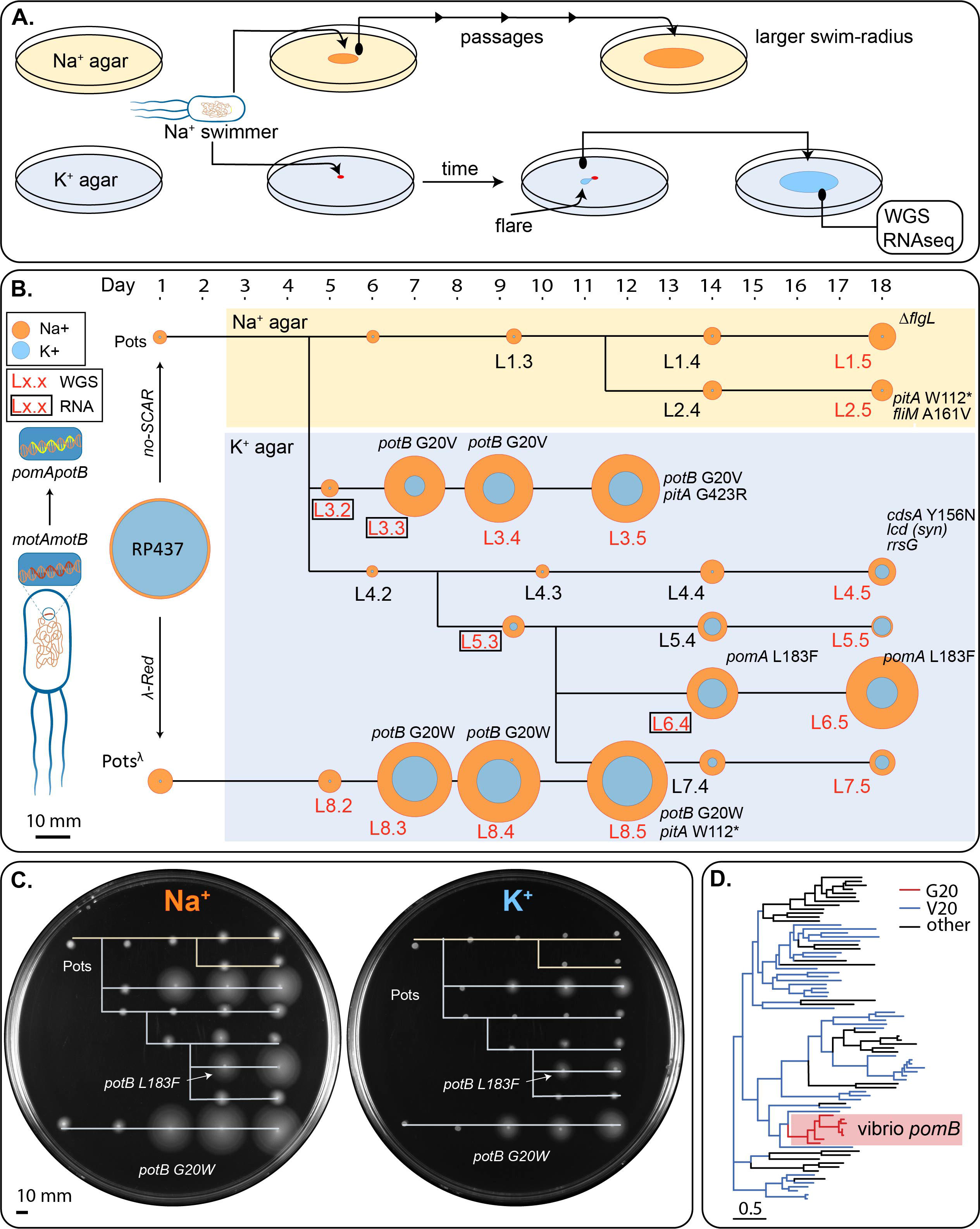
Directed Evolution of the Flagellar Motor. A) Experimental overview. Sodium swimming strain is repeatedly passaged on either Na^+^LB (~100 mM [Na+]) or K^+^LB (~15 mM [Na+]) plates. Where flares are observed on potassium plates, indicating a potentially upmotile variant, these are propagated and sent for sequencing. B) Our edited *E. coli* strains Pots and Pots^λ^, obtained from *E. coli* RP437 via no-SCAR and λ-Red recombineering respectively, were passaged on soft agar (colored background, yellow: Na+; blue: K+) over an 18-day period. The summary comprises a total of 8 lineages (L1-L8) selected for further investigation, each comprised of 5 members (ie. L1.3 indicates the first lineage and the third passage of a motile flare). The day of collection and re-inoculation is indicated above each lineage member. The ability of each lineage member to swim in the presence of high or low sodium is displayed by a yellow or blue ring, respectively, corresponding to swim size on a swim plate. Lack of motility after 24 hr of incubation on K^+^ soft agar is represented by a blue dot which corresponds to colony growth only. Ring sizes were measured from the pair of 150 mm diameter swim plates presented in panel C. Colonies that were non-motile in K^+^LB swim plates were confirmed with further incubation (Supplementary Fig. 3C), and are indicated by blue dots at innoculation centre. Lineage labels are red to indicate whole genome sequencing (WGS) data availability and boxed to indicate RNAseq data availability. Ancestral Pots was also analysed by WGS and RNAseq for comparison. SNPs identified via variant calling relative to the Pots reference genome are annotated next to each respective lineage member. Highlighted genes other than *pomA* and *potB*: *pitA* (metal phosphate:H^+^ symporter), *flgL* (flagellar hook-filament junction protein 2), *fliM* (flagellar motor switch protein), *cdsA* (cardiolipin-diglyceride synthase), *icd* (isocitrate dehydrogenase), *rrsG* (16S ribosomal RNA). Scale bar: 10 mm. A full list of identified SNPs is provided in Supplementary Table 1. C) Recapitulation of the directed evolution experiment. Na^+^ (left) and K^+^(right) soft agar plates inoculated with a 1 μL aliquot of glycerol stock of each strain indicated in (B (except RP437) and arranged in the same order as (B. Arrows and labels indicate the first member in each lineage to display a mutation in the stator genes PomA or PotB. Each plate was incubated for 24 hr at 30°C. D) Phylogeny of MotB across 82 species with ancestral reconstruction at the G20 site. G20 is conserved in the *Vibrio spp.* clade. Full phlogeny is shown in Supplementary Fig. 14.

### Whole genome sequencing of evolved lineages

Lineages were selected for whole-genome sequencing (WGS) after a preliminary screening for mutations in the stator genes by Sanger-sequencing PCR amplicons spanning the genomic *pomApotB* locus (Fig. 1). Variant calling to the MG1655 reference genome was used to compare single nucleotide polymorphisms (SNPs) between members of the same lineage. Our intended *pomApotB* edit was the only difference between the RP437 and Pots genomes, indicating that neither no-SCAR nor λ-Red editing had resulted in off-target edits. 153 SNPs were called as variants between our experimental parent RP437 and the MG1655 reference which were shared across all lineage members (Supplementary Table 1).

Several lineages whose descendants could swim in reduced sodium had mutations at the *pomApotB* locus (L3.3-4-5: *potB* G20V; L6.4-5: *pomA* L183F; L8.3-4-5: *potB* G20W). In contrast, lineages passaged only on ~100 mM Na^+^ agar (L1 & L2) accumulated mutations not in stators but in the flagellar components. Lineages passaged on ~15 mM sodium-poor agar whose descendants could not swim (L4) had no mutations on any flagellar genes.

### Differential expression across upmotile lineages G20V and L183F

To determine which genes may be involved in adaptation, we performed RNAseq experiments to measure transcript levels for two lineages which evolved different stator mutations over different lengths of time (Fig. 2, Supplementary Table 2). These were: *pomA* L183F (Na^+^ powered phenotype, 14 days) and *potB* G20V (H^+^ phenotype, 7 days). From these, we selected the common ancestor to the two lineages (Pots), the last lineage member before the mutation occurred (L5.3 and L3.2, the pre-fixation sample) and the first available member carrying that lineage mutation on the chromosome (L6.4 and L3.3, the fixed variant). Of note, L5.3 displayed improved swimming in K^+^LB despite being isogenic to its parent strain, the Pots common ancestor. Member L3.2 of the G20V lineage, the immediate predecessor to L3.3, in contrast, was non-motile. We closely examined the transcripts mapped to the *pomA/potB* locus in all samples and saw no enrichment of mutant transcripts above the background noise in the pre-fixation samples (Supplementary Fig. 8). This confirmed that a chromosomal mutation was responsible for the observed stator variants. To identify the common processes which could lead to variant fixation we calculated which differentially expressed genes were present in both lineages and performed pathway analysis to identify which biological processes were relevant to the shared genes (Fig. 2).

**Fig.2.**
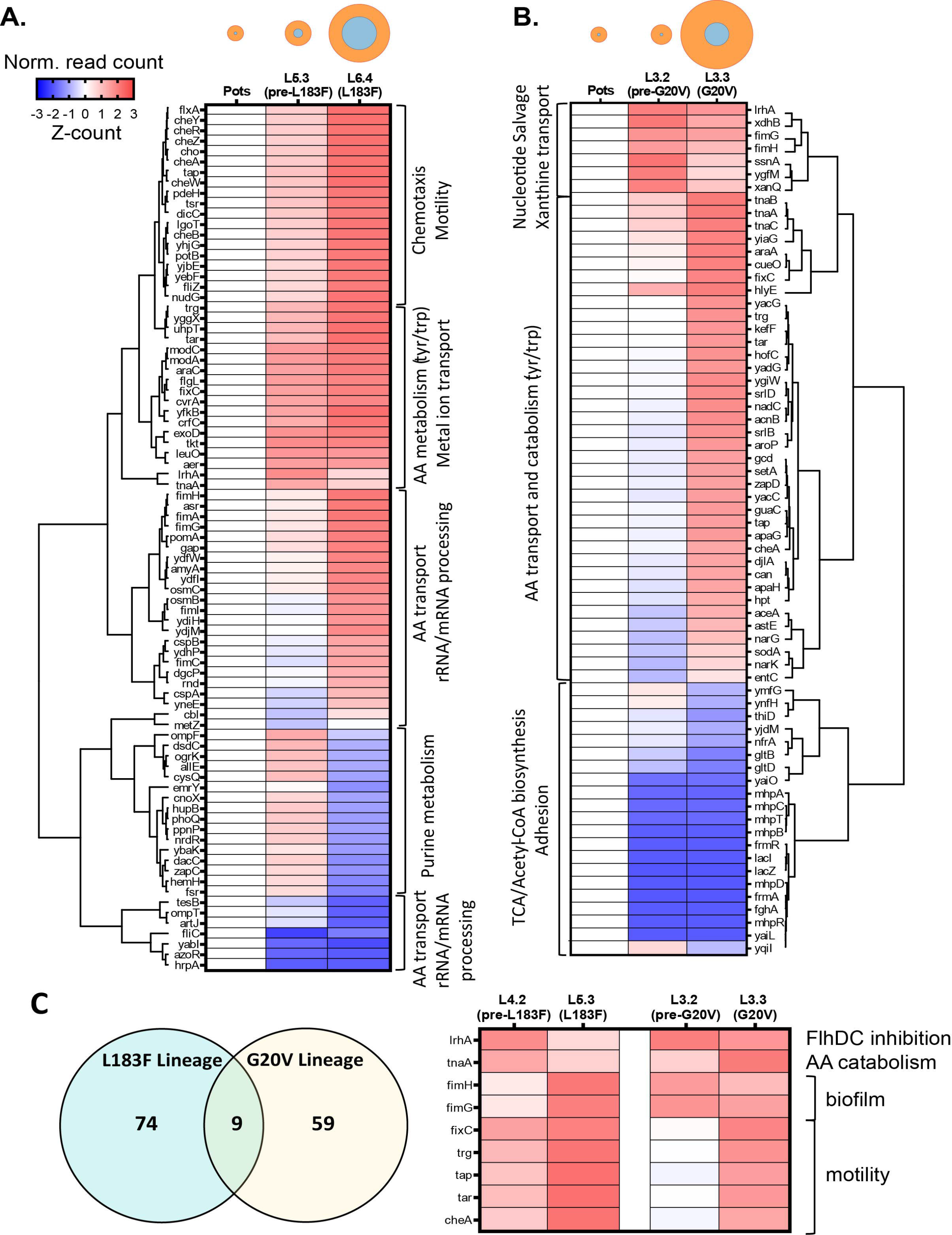
Differential expression analysis of RNAseq data. Differentially expressed genes (DEGs, adjusted p-value < 0.01) measured across selected members of the L183F lineage (A) or G20V lineage (B). The average read counts (n = 3) for each gene were normalized by z-score and displayed as clustered heatmaps, flanked by their respective dendrograms. DEG clusters are labelled by their representative biological process (PANTHER). Genes upregulated with respect to the Pots strain are labelled in red, downregulated ones in blue. The ring diagrams above each heatmap are taken from fig.1 and indicate the motility phenotype of each lineage member. C) Venn diagram indicating the number of DEGs in each dataset and the DEGs in common, shown in the heatmap on the right.

Both lineages increased expression of *lrhA* at the pre-fixation point, which acts as an inhibitor of the *flhDC* flagellar master regulator. Expression response was dominated by pathways under control of RpoD (σ70) / FliA (σ28) in L183F, and RpoD (σ70) in *potB* G20V (Supplementary Fig. 9). The prominence of *FliA* regulation in *pomA* L183F was reflected by upregulation of motility genes at the fixation stage. Those same genes were not significantly affected in the G20V lineage, which instead promoted primarily adhesion/biofilm behavior in response to the changed environment.

### Phenotypic characterization of evolved strains in the presence and absence of sodium

We characterized rotational and free-swimming phenotypes of the parent and evolved strains in the presence and absence of sodium using a tethered cell assay with particular focus on the upmotile G20V variant, L3.3 (Fig. 3, Supplementary Fig. 10). The *potB* G20V fixed variant was clearly distinguishable from Pots in this assay and maintained rotation in both the absence of sodium and in the presence of phenamil, an amiloride derivative and sodium-channel blocker (Fig. 3A) (*22*). We confirmed the *potB* G20V H^+^-powered phenotype by introducing the same point mutation (GGG to GTG) on our plasmid vector (pPots) and testing motility in tethered cell assay when the protein was expressed in the statorless Δ*motAB* RP6894 (Fig. 3B). Measurement of tethered rotational speed vs lithium and vs sodium showed that the *potB* G20V variant was motile at 0 mM Na^+^ and 0 mM Li^+^, with a dependence on the concentration of ion, in contrast to *motA/motB*, whose rotational speed was independent of sodium or lithium concentration (Supplementary Fig. 11A-C). All stator types had cessation of rotation at 50 μM CCCP and showed no measurable effect as pH was varied between 6.0 and 8.0 (Supplementary Fig. 11DE).

**Figure 3.**
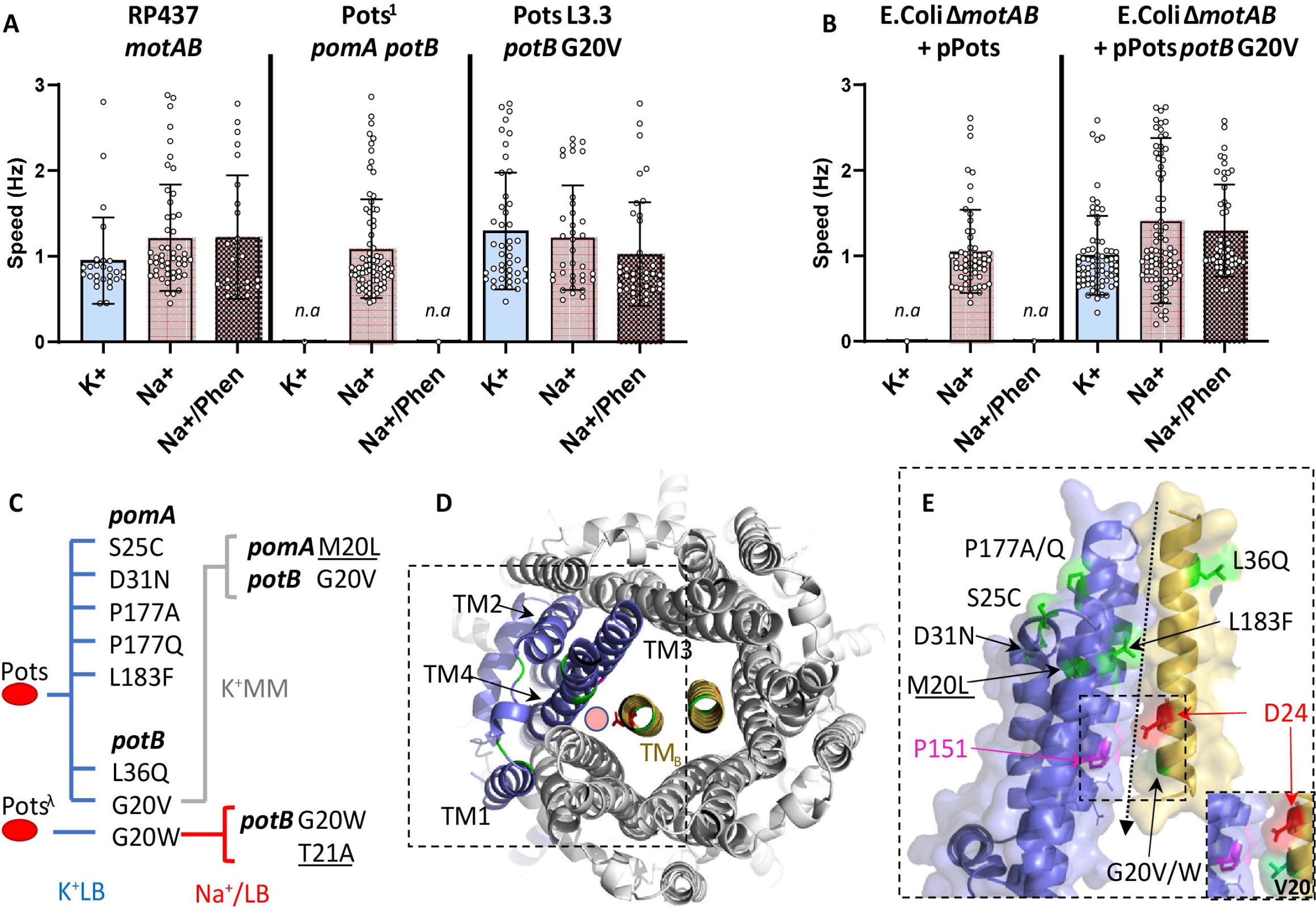
Functional characterization of evolved stators. A) Single cell speed measurements using the tethered cell assay measured in Hz (revolutions/s). Blue bar indicates speed in K^+^ motility buffer (K^+^MB), red bar: Na^+^ motility buffer (Na^+^MB); red patterned bar: Na^+^MB + 100 μM phenamil. Number of cells analysed per condition (from left to right): RP437: 27, 51, 25; Pots: n.a, 78, n.a; Pots L3.3 *potB* G20V 45, 36, 39 (n.a. indicates no visible rotating cell). Error bars indicate Standard Deviation (S.D.). B) Single cell speed measurements using the tethered cell assay in RP6894 Δ*motAB* strain co-expressing *pomA* and *potB* G20V from pPots plasmid. Blue bar indicates speed in K^+^MB, red bar: Na^+^MB; red patterned bar: Na^+^MB + 100 μM phenamil. Number of cells analysed per condition (from left to right): (Δ*motAB* + pPots: n.a., 32, n.a; Δ*motAB* + pPots *potB* G20V: 40, 63, 48). Error bars indicate S.D. Single cell tracked data shown in Supplementary Fig. 12. C) Graphical summary of stator gene mutations detected across all directed evolution experiments and the growth conditions under which these mutations arose. LB indicates agar containing yeast extract and tryptone. MM indicates agar in minimal media. Mutations in a subsequent generation are underlined. D) View from the extracellular side of the transmembrane portion of PomA_5_PotB_2_ stator complex (see Methods: Structural Modelling). One monomer of subunit A is coloured in blue and the TM domains of the B subunits are coloured in yellow. Mutant sites obtained in the directed evolution experiments are labelled in green, the catalytic aspartate residue essential for function is highlighted in red. The red circle indicates the predicted location of the ion transport pore (inward conduction). E) side view of the area highlighted by the dashed box in D. Residues P151 (PomA) and D24 (PotB) are also highlighted in magenta and red respectively. Primary mutation sites are indicated in black, while secondary mutation sites are underscored. The arrow at the interface between (A)TM3-4 and (B)TM indicates the predicted location of the ion transport pore. The inset highlights the change in the pore region due to the G20V substitution in PotB (green).

We further tracked single cell rotation in the tethered cell assay with sequential exchange of buffers, including characterization of *pomA* P177Q and *pomA* L183F (Supplementary Fig. 12). There, *potB* G20V (L3.3) actively rotated in K^+^MB_EDTA_ and in Na^+^MB + 100 μM phenamil. K^+^MB cont indicating sodium-independent rotation. We further synthesized all possible variants at site G20 (*23*), and, of these, only G20V was able to rotate in the absence of sodium (Supplementary Fig. 13).

### Natural prevalence of *motB* G20 and *motB* V20

We examined the natural prevalence of glycine and valine at site 20 across a phylogeny from a collection of 82 *motB* sequences. Of these sequences, G20 was present only in the clade corresponding to *Vibrio sp. pomB* (7 sequences), whereas V20 was distributed across the phylogeny more broadly (41 sequences) (Fig.1D, Supplementary Fig. 14). Ancestral reconstruction across all nodes predicted that the ancestral phenotype of this phylogeny was V20.

### Reproducibility of stator mutagenesis and capacity for reversion

To examine mutation reproducibility in the stators, we subjected 55 Pots colonies to directed evolution in K^+^LB swim plates. These yielded a total of 42 flares within the first 3 days of incubation, which were then passaged four more times at 3-day intervals. At the end of this experiment we selected the 20 terminal lineage members which produced the largest swim rings and Sanger-sequenced them at the *pomApotB* locus. For these we observed a total of 5 mutations in *pomA* (S25C, D31N, P177A, P177Q and L183F) and one more new mutation in *potB* (L36Q) (Fig. 3C). We mapped monomer models for PomA and PotB generated using Alphafold (*24*) to a model PomA_5_PotB_2_ complex using the published high-resolution *B. subtilis* MotA_5_B_2_ structure (*10*) (Fig. 2C & 2D). All stator mutations accumulated at sites proximal to or within the predicted ion-transport pore, at the interface between the PotB transmembrane domain and the third and fourth transmembrane domains of PomA (Fig. 2D).

Over the course of these experiments, we found that three stator residues underwent mutation at the same site twice (PomA L183F (2x), PomA P177A & P177Q, PotB G20V & G20W). WGS revealed that the *pitA* gene had mutated in three separate lineages (L2.5, L3.5, L8.5) with one of the mutations occurring twice (*pitA* W112*Stop).

To test for the capacity for reversion, we took sequenced lineages that swam in K^+^LB swim plates (L3.3-4-5, L6.4-5, L8.3-4-5) and reintroduced them to an environment with high sodium (Na^+^LB swim plates) (Supplementary Fig. 15A). After 10 rounds of daily passaging, no reversion in the mutants that had enabled low-sodium swimming was observed, with only a single additional mutation gained: a *potB* T21A mutation in the terminal descendant of *potB* G20W *pitA* W112*stop (L8.5).

### Directed evolution from later starting points

We tested whether evolution could be more easily directed on minimal media when starting from a more favorable vantage point. We examined all stator mutants that swam on K^+^LB swim plates in conditions of further sodium scarcity (K^+^MM: ~1 mM [Na^+^]) (Supplementary Fig. 15B, Supplementary Table 3). Initially, only the strains with *potB* G20V and *pomA* P177Q mutations could swim, and this capacity was maintained following five passages over 12 days. Sanger sequencing revealed that the terminal descendant of p*otB* G20V *pitA* G432R mutant (L3.5) gained a further mutation in *pomA* (M20L). A summary of all stator gene mutations obtained from all directed evolution experiments is provided in Fig. 3C. We also verified the capacity of strains at the pre-mutation stage to replicate the mutation of their lineage and found that L3.2 (pre-G20V) consistently replicated the *potB* G20V mutation over two independent experiments (6 out of 6 flares), and in one instance incorporated a *potB* L36Q mutation in addition to G20V (Supplementary Fig. 16).

### Comparison of fitness and motility in competition

Finally, we compared fitness and competitive motility directly between the L3.3 (*potB* G20V) and the Pots ancestor. The *potB* G20V variant conferred a clear and discernible motility advantage in a mixed population on K^+^LB swim plates, while displaying similar growth in both Na^+^ and K^+^ LB liquid media (Supplementary Fig. 17).

## DISCUSSION

Mutation in DNA is a critical requirement for adaptation and evolution. Much is known about sources of spontaneous mutagenesis in bacteria, but the regulatory and molecular processes which control adaptation are not so well understood (*25, 26*). Epistasis and functional redundancy in biochemical pathways impede the accurate forecasting of mutagenic events responsible for rescuing a phenotype which could be the result of loss-of-function mutations in negative regulator or gain-of-function in positive regulators (*27*). The flagellar system and its regulatory elements have been known to be under selective pressure due to their associated energetic and fitness costs, not always resulting in positive selection but also often resulting in gene deletion or inactivation by insertion sequences (*28–33*). By targeting the stator subunit of the flagellar motor, we have been able to study the molecular events leading to the spontaneous adaptation of a unique module within a highly conserved molecular complex (*34*).

Previous reports have shown that bacterial motility can adapt (*8*) and be rescued (*35*) via remodelling of the flagellar regulatory network. Ni et al. observed that evolutionary adaptation of motility occurs via remodeling of the checkpoint regulating flagellar gene expression (*8*). Their experiments tracked adaptive changes in swim plates, matching our experiments, but their only selection criteria were for improved swimming in an unhindered swimming population. In agreement with our results (*fliM* A161V), they found *fliM (*M67I and T192N) to be the amongst the first genes to mutate in the improved swimmer population but they did not report any changes in *flgL* (ΔA57-Q58), nor, significantly, did they see any mutations in any stator genes. Flagellum-mediated motility also appears to be naturally robust to the loss of regulatory factors, such as the enhancer-binding protein fleQ in *P. fluorescens*, which function can be substituted by distantly related homologous proteins (*35*).

In contrast, our *E. coli* Pots strain faced selective pressure from ion scarcity. Our scenario is reminiscent of previous semi-solid agar experimental evolution studies on the adaptation of antibiotic resistance and produced similar results. In the MEGA plate experiments of Baym et al., they similarly saw that the phosphate transporter *pitA* was repeatedly mutated, often to a frameshifted or nonsense variant (*21*). In similar experiments the isocitrate dehydrogenase *icd* was also seen to mutate often (*1, 36*).

Differential expression analysis of our RNAseq datasets revealed that our two chosen lineages displayed different transcriptomic signatures during the adaptation of the stator genes. While the L183F lineage displayed many upregulated genes involved in motility, the G20V lineage was found to regulate genes involved in biofilm formation.

The improvement in low sodium motility observed in L5.3 (pre-L183F mutation stage) in comparison with its Pots ancestor may be explained by the upregulation of flagellar and chemotaxis genes in L5.3. Notably, L6.4 (pomA L183F) did not swim in the total absence of sodium (Supplementary Figure 12D), however the swim ring size of both L5.3 and L6.4 were greater relative to Pots on low sodium K^+^LB plates. The genes involved have also been measured as upregulated in a similar study on the adaptation of *E. coli* to swimming in soft agar (*8*) and it could be the case that upregulation of several flagellar components, including the chimeric sodium stators themselves (PomA and PotB), improves motility in these ~15 mM [Na^+^] plates. In contrast, the expression profile of the Na^+^-dependent swimmer L3.2 (pre-G20V mutation stage) was characterized by the regulation of genes involved in metabolic pathways indicative of nitrogen starvation, fermentation of products of catabolism such as amino acids and nucleotides and the transition to a biofilm lifestyle. Roughly 10% of genes showed significant changes in transcription in both lineage trios (10.8% and 13.2% in L183F and G20V respectively) and all of these were upregulated during adaptation.

After the G20V mutation was fixed, L3.3 (G20V) was found to upregulate chemotaxis and motility genes, and other markers of adaptation to soft agar (*cheA*, *trp*, *tar*, *tap*) (*8*). Both lineages upregulated the *flhDC* regulator *LrhA* at the pre-mutation stage, hinting that downregulation of the flagellar biosynthetic cascade is a shared trait in the early stages of adaptation to a sodium-poor environment. Similarly, both lineages upregulated nucleotide catabolic processes and salvage pathways, a feature also observed in antibiotic response and which can affect mutation rates by disturbing the NTP balance in the cellular pool (*37–39*). This might suggest that the mutations were a product of stress-induced mutagenesis, a known facilitator of evolution (*40*) which has been proposed to involve *RpoS*-mediated upregulation of the *DinB* error-prone polymerase (*41*). In our RNAseq data, we saw minimal involvement of *RpoS* signalling (Supplementary Fig. 9) and no evidence of upregulation of known error-prone polymerases (Supplementary Table 2). This suggests that the molecular events leading to fixation are not resolvable from transcriptomic analysis of only two lineages, or that alternate mechanisms could have facilitated mutagenesis. These may include mutagenesis via redox events on DNA, as seen in antibiotic resistance (*42*), or via transcription-dependendent mechanisms (*43, 44*). The biological pathways leading to mutation remain to be elucidated.

We observed a convergence of mutations on the stator genes and even to the very same nucleotide (GA<u>G</u> (L) to GA<u>A</u> (F) in two separate *pom*A L183F lineages. Stator genes were the first to mutate in all of our WGS lineages under pressure from sodium scarcity. Given this was from a clonal population under identical environmental constraints, it suggests that adaptation of the stators provides a strong selective benefit in changing environments.

Mutations in stators are known to affect ion usage and may confer dual ion coupling capacity (*12–14, 45*). For example, the substate preference of the *B. alcalophilus* MotPS stator (Na^+^/K^+^ and Rb^+^) was changed with the single mutation M33L in *motS*, causing the loss of both K^+^- and Rb^+^-coupling motility in *E. coli* (*46*). Similarly, a bi-functional *B. clausii* MotAB stator (Na^+^/H^+^) triple mutant (*motB* V37L, A40S and G42S) was selective only for sodium ions while the combination of mutations G42S, Q43S and Q46A made MotB selective only for H^+^ (*47*). The previously reported S25C (*48*) and D31N (*49*) amino acid substitutions in PomA have been shown to reduce motility and, in the case of D31N, affect ion usage. Furthermore, single point mutations in stator genes of *Vibrio* spp. (eg. *pomB* G20V / G20R / P16S) have been shown to impart phenamil resistance both in *Vibrio alginolyticus* and in our Pots strain (*50–52*).

The PotB-G20V variant directly evolved here is capable of rotation in the absence of sodium, but its swimming speed does increase with increased availability of either sodium or lithium, with an energisation profile more akin to the parental Pots strain than the proton powered *E. coli* wild-type (Supplementary Fig. 11). This perhaps suggests that PotB-G20V is an opportunistic stator adaptation that enables promiscuity, allowing the passage of protons in sodium-scarce conditions.

The key difference in this work compared with previous efforts for directed evolution of the stators via mutagenesis (*53*), (*54*) is that here we edited the stators directly onto the *E. coli* genome to direct stator evolution *in vivo* in the native *E. coli* genomic context. This would not be possible in *Vibrio sp.* since *Vibrio* cells do not survive at low sodium. Conversely, in our system it is difficult to use directed evolution to revert the ion-selectivity of the stator (H^+^ to Na^+^) because in *E. coli* it is not possible to drastically reduce the proton concentration without affecting essential systems. Nevertheless, the observation of no revertant agrees with previous work suggesting that requirements for Na^+^ binding are more strict than for H^+^ binding, and that mutations that convert a Na^+^ motor to an H^+^ are more accessible than the reverse (*55, 56*).

Mutation of pore-proximal residues into hydrophobic residues (eg. G20V) hinted at a mechanism for varying constrictions in the pore to alter the efficiency of ion binding. However, in contrast, none of the bulkier, hydrophobic amino acid replacements at *potB* G20 (eg. F,W) resulted in a similar G20V-like H^+^-powered phenotype (Supplementary Fig. 13C). This suggests a selectivity mechanism enabled by G20V that is not driven simply by size. We propose an alternate mechanism whereby selectivity is maintained through perturbing the electrostatic environment in the vicinity of PotB D24 and the conserved P151 of PomA (*E. coli* MotA P173) (*57*).

Upon examination of the phylogenetic record with specific focus on the G20V locus, we observed that valine was more prevalent and distributed more broadly across microbial strains. This contemporary prevalence, and ancestral sequence reconstruction across our phylogeny implied that the ancestral state of MotB was more to be likely V20. While G20V point mutations arose spontaneously in our experiments within a few days, these transitions do not appear to have occurred in the evolutionary record. This may indicate constraints on the adaptation of the sodium-powered stator units when considered in their native sodium-dependent hosts.

Caution is required when applying learnings from directed evolution to natural evolution since selection pressures in the wild are not typically general (*58*). In this study we have leveraged our system, and our experimental design, to obtain large, measurable phenotypic change through a single mutation at the G20 locus. The stators of the flagellar motor appear ready to evolve in our experiments: they do not require transit through additional cryptic or neutral mutations, and thus are a model system for exploring molecular adaptation of ion selectivity. Alternative candidates such as sodium porters and pumps often have built in redundancy (*59*), and many marine microbes require sodium to be viable and thus evolutionary pressure cannot be applied with such specificity to a single protein complex. In our idealised system, we are able to examine isogenic bacterial colonies without competitive effects, in a homogeneous medium without local niches. Nevertheless, it still remains difficult to quantify rates of adaptation (*60*). For these reasons, our system is optimised to produce stator variants, but it may be that this ease for adaptation via single-point mutation is accessible precisely because cryptic mutations have been accumulated due to exposure to changing environments in the distant past (*61, 62*).

Motility confers a fitness advantage that is worth significant energetic investment despite the high cost of synthesizing the flagellar machinery (*4, 63*)(*33*). This advantage can only be seized if the correct ions are available for stator-conversion into torque. We have shown here that the flagellar apparatus is capable of single-site mutation to adapt the stator genes to harness other available ions. While the modularity of the overall flagellar motor is now well shown (*64*) here we have observed further modularity and adaptability within the stator complex. Chimeric functional stators can not only be engineered from stator components in various species, they are also subsequently capable of rapid mutation to use ions more promiscuously. Motility provides many benefits to an organism but the exact evolutionary event which resulted in the first flagellar motor is not known (*65*). Nevertheless, our work shows that, following emergence, subsequent environmental adaptation can occur rapidly.

## MATERIALS AND METHODS

### *E. coli* strains, plasmids and culture media

*E. coli* strain RP437 was used as the parent strain for genomic editing experiments (*66*), (*67*). The pSHU1234 (pPots) plasmid encoding *pomA* and *potB* (*50*) was used as the template to generate the double stranded donor DNA. This was used to replace the *motA* and *motB* gene on the RP437 chromosome. Point mutations in plasmids were generated using the QuikChange™ technique while saturation mutagenesis of the *potB* G20 site was performed using the ‘22-c trick’ technique (*23*). Liquid cell culturing was done using LB broth (NaCl or KCl, 0.5% yeast extract, 1% Bacto tryptone). Cells were cultured on agar plates composed of LB Broth and 1% Bacto agar (BD, U.S.A.). Swim plate cultures were performed on the same substrates adjusted for agar content (0.3% Bacto Agar). Minimal Media (MM) was used to replace Yeast Extract and Tryptone in soft agar swim plates. MM composition: 10 mM KH_2_PO_4_ (KPi), 1 mM (NH_4_)_2_SO_4_, 1 mM MgSO_4_, 1 μg/ml Thiamine, 0.1 mM of each of the amino acids Thr, Leu, His, Met and Ser. Inhibition of Na^+^ -dependent motility was performed using phenamil methanesulfonate (P203 Sigma-Aldritch) at 50 μM and 100 μM concentrations while H^+^-dependent motility was inhibited using the protonophore Carbonyl cyanide 3-chlorophenylhydrazone (CCCP, C2759 Sigma-Aldritch).

### Editing *E. coli* with Cas9-assisted Recombineering

This procedure was adapted from the no-SCAR method (*19*). The target strain to be edited (*E. coli* RP437) was sequentially transformed first with the pKD-sgRNA-3’MotA (Sm^+^) plasmid, encoding a sgRNA sequence directed at the 3’ end of *motA*, and then with the pCas9cr4 (Cm^+^) plasmid to yield a parent strain harboring both plasmids. The overlap extension PCR technique (*68*) was employed to assemble linear double stranded DNA molecules (dsDNA) using 3 starting dsDNA fragments. The resulting donor DNA was electroporated in the plasmid-bearing host and the successfully edited clones were selected via colony PCR and Sanger sequencing of the *motAB* locus. A list of primers and PCR protocols used in this work is provided in Supplementary Fig. 18.

### Construction of Pots by λ-Red Recombineering

Chromosomal replacement from *motAmotB* to *pomApotB* was achieved by using a λ-Red recombination system, with plasmid pKD46 encoding the Red system and positive selection for the recovery of swimming ability (*69*). Motile clones were selected by isolating motile flares on swim plates.

### Measurement of sodium concentration in solutionsusing Atomic Absorption Spectroscopy (AAS)

The amount of residual sodium in motility buffers and culture media was measured using Atomic Absorption Spectroscopy (ANA-182, Tokyo Photo Electric co., LTD, Japan). AAS measurements are displayed in Supplementary Table 3.

### Tethered cell assay preparation and analysis

The tethered cell assay with anti-FliC-antibody (*69, 70*) was performed as previously described (*22*). Briefly, 1 mL of cells (OD_600_=0.5) grown in K^+^TB buffer (Tryptone, 85 mM KCl) was sheared by passing the cells suspension through a 26-gauge syringe needle 30 times. These cells were then washed 3 times in 1 mL motility buffer (K^+^MB: 85 mM KCl, 10 mM KPi, pH=7.0) and finally resuspended in 500 μL of motility buffer. Then, 20 μL of suspension was loaded into a tunnel slide pre-filled with motility buffer that had previously been incubated with anti-FliC antibodies for 15 min at room temperature (1:300 dilution in water). The unbound cells were then removed from the tunnel slide by washing with a total of 200 μL of motility buffer (~10 times the tunnel slide volume). The tethered cells time lapse videos were recorded at 40x magnification on a phase contrast microscope (Nikon). Time lapse videos were collected using a camera (Chameleon3 CM3, Point Grey Research) recording 20s-long videos at 20 frames per second. Experiments involving single cell tracking during buffer exchange were recorded at 60 frames per second, with cells washed and resuspended in K^+^MB + 0.1 mM EDTA-2K (K^+^MB_EDTA_). Free-swimming cells were grown overnight in K^+^TB at 30°C to OD_600_ ~0.5 then washed 3 times in 1 mL K^+^MB before resuspension in 500 μL of K^+^MB and imaging in a tunnel slide. A custom LabView software (*17, 22, 50*) was employed as previously reported to estimate specific rotational parameters of the tethered cells such as rotation frequency (speed), clockwise and counterclockwise bias, switching frequency and speed of swimming cells. FliC-sticky *E. coli* RP437 cells (Δ*motAB* Δ*cheY* Δ*pilA fliC*^st^) (*71*) were used to collect data presented in Supplementary Fig. 11. Visualization of the data was performed using Graph Pad Prism 8.

### Fitness comparison assay between Pots and L3.3 (*potB* G20V)

Overnight cultures of the two strains grown in K^+^LB were adjusted to equal OD_600_ then mixed at 1:1, 1:10 and 1:100 (L3.3 : Pots) ratios in a 100 μl volume. 10 μl of each mixture was then used to inoculate a 2 ml K^+^LB liquid culture at 30°C for 24hr. The resulting dense culture was then diluted to OD_600_=0.25 and then further diluted 10^6^-fold in K^+^LB before streaking onto K^+^LB swim plates (20 μl streaks) and incubating for 48hr at 30°C.

### Single Nucleotide Polymorphism (SNP) analysis

Whole genome sequencing of 22 *E. coli* strains was performed using a MiSeq 2 x 150bp chip on an Illumina sequencing platform. Sequencing was carried out at the Ramaciotti Centre for Genomics, Kensington and delivered as demultiplexed fastQ files (Quality Control: >80% bases higher than Q30 at 2×150 bp). The SNP calling and analysis was performed using Snippy (*72, 73*). The short reads from sequencing were aligned to the MG1655 reference *E. coli* genome (GenBank: U00096.2) and to a synthetic genome based on MG1655, edited to contain the Pots stator sequences from pPots (*pomA*/*potB*) at the *motAB* locus.

### Transcriptomics

RNA was extracted from bacterial cultures inoculated with glycerol stocks of the relevant strains and grown in K^+^LB broth at 30°C until OD_600_=0.5. Total RNA was extracted from a 0.5 ml aliquot of the culture using the RNAeasy Protect Bacteria Mini Kit (74524, QIAGEN) with on-column DNAse digestion, as indicated in the manufacturer protocol. RNA quality was assessed using a TapeStation System (RNA ScreenTape, Agilent). All RNA samples selected for sequencing had an RNA Integrity Number (RIN) > 8. Library preparation and sequencing were performed at the Ramaciotti Centre using the NextSeq 500 platform (Illumina) running for 150 cycle using a MID flowcell in paired-end read mode (2×75bp). Fastq files containing the RNAseq reads underwent quality control using FastQC (*74*) and then processed with FASTP (Version 0.20.1)(*75*), to remove low quality reads and trim adaptor sequences. Reads were aligned to the Pots reference genome using HISAT2(*76*), transcripts were assembled and quantified using STRINGTIE (*73*) and differential expression analysis was carried out using DESeq2 (*77*). Heatmaps dendrograms were generated using the Heatmap2 function from the R gplots package (Heatmap2). Complete clustering was performed using the Euclidean Distance method. All the analysis tools described above were run on the Galaxy webserver (https://usegalaxy.org/). Nucleotide variations that were present in motB and pomA were quantified from RNA-Seq data using the Rsamtools pileup function (*78*). This involved the writing of custom R scripts (available at https://github.com/VCCRI) that compared Rsamtools output to the genome reference.

We then performed pathway analysis to identify which biological processes were relevant to the shared genes using the EcoCyc database (*79*).

### Structural Modelling

The PomA_5_PotB_2_ model was assembled by modelling each monomer using the Colabfold pipeline (*24*), and by aligning the resulting monomers to each subunit of the *B. subtilis* MotA_5_B_2_ structure (PDB:6YSL) (*10*).

### Phylogenetics and Ancestral Reconstruction

Phylogeny was generated with RAxML-HPC v.8 on XSEDE (*80*) through the CIPRES Science Gateway (*81*). The phylogeny was calculated using the PROTGAMMA protein substitution model, LG protein substitution matrix, and a parsimony seed value of 12345. Ancestral sequences were calculated using CodeML, a maximum likelihood program from the PAML package, using the LG rate file with the Empirical+F model and using 8 categories in dG of NSsites models (*82*). Ancestral Sequence Reconstructions at each node were used to determine G20/V20 identity at each node. Genomic context for the stators was was pulled from the KEGG Database (*83*).

## Supporting information

List of WGS SNPs

## AUTHOR CONTRIBUTIONS

PR and MABB designed experiments and executed experiments in strain editing, molecular biology and microbiology. TI and YS executed experiments in strain editing. MABB executed bioinformatics surrounding variant calling. AL and MABB executed bioinformatics surrounding phylogenetics. PR, DTH, EG and MB executed bioinformatics surrounding the transcriptomics. MB supervised the design, execution and writing of the project. All authors contributed to writing and revision of the manuscript.

## ACKNOWLEDGEMENTS

We would like to acknowledge Myu Yoshida and Rie Ito for technical assistance.

## COMPETING INTERESTS

The authors declare that they have no competing interests.

## DATA AVAILABILITY

All data needed to evaluate the conclusions in the paper are present in the paper and the Supplementary Materials. All WGS and RNAseq data is deposited as Bioproject Accession: PRJNA729860. Pots genome is deposited as GenBank: CP083410.1.

## FUNDING

YS was supported by JSPS KAKENHI (JP18H02475 and JP20K06564), MEXT KAKENHI (JP19H05404, JP 21H00410 and JP21H05892) and Takeda Science Foundation. MABB was supported by a UNSW Scientia Research Fellowship, ARC Discovery Project DP190100497 and HFSP Young Investigator Project Grant RGY0072/20.

## Notes

### Competing Interest Statement

The authors have declared no competing interest.

### Summary of Updates

Updated Fig. 1, added text to results/discussion, added additional SI Figures.

