## Supplementary Figures 1-18, Supplementary Tables 1-3 for "The rapid evolution of flagellar ion-selectivity in experimental populations of *E. coli*"

**Supplementary Information**

**Supplementary Figures**

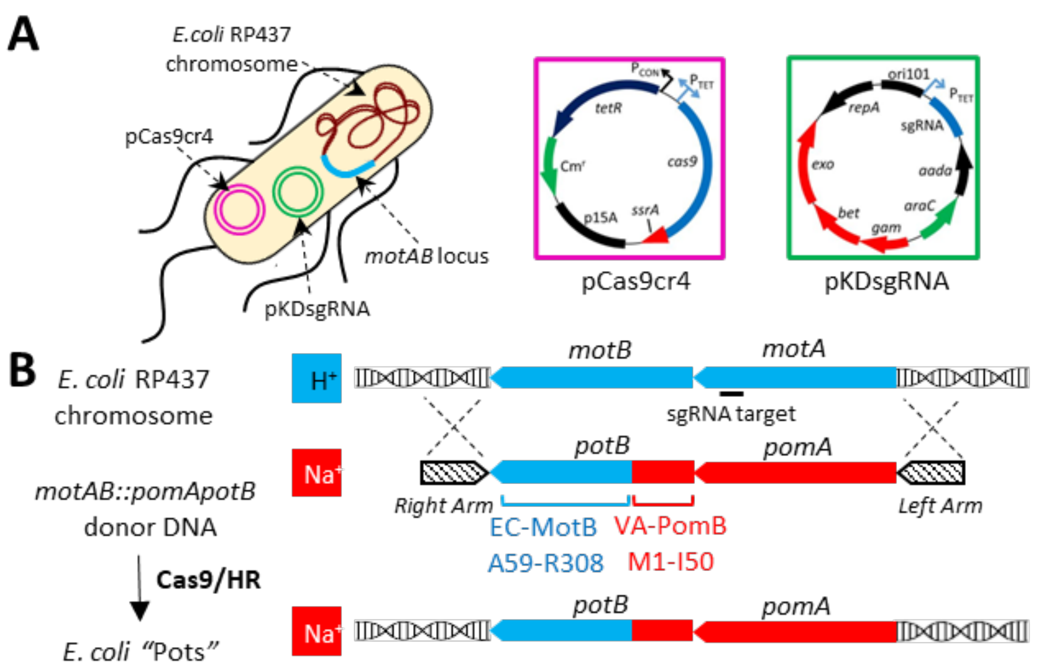


**Supplementary Fig. 1. Genome editing of E. coli RP437 using the no-SCAR method.** A) Schematic of an *E. coli* RP437 cell carrying the pCas9cr4 (pink) and pKDsgRNA (green) plasmids required for no-SCAR recombineering at the *motAB* locus (cyan) on the bacterial chromosome. The map of each plasmid is provided in the colored boxes. B) The *motAB* locus encoding the $H^+$ powered *E. coli* stator proteins is replaced with *pomApotB* encoding a chimeric $Na^+$-powered stator by using a linear double-stranded DNA donor template for homologous recombination (HR). This is flanked by a left and right homology arm to facilitate HR at the *motAB* locus after Cas9 has cut the chromosome at its sgRNA target near the 3' end of *motA*. *potB* is composed of *V. alginolyticus pomB* Met1-Ile50 and *E. coli motB* Ala59-R308. The resulting $Na^+$-powered *E. coli* strain was renamed "Pots".

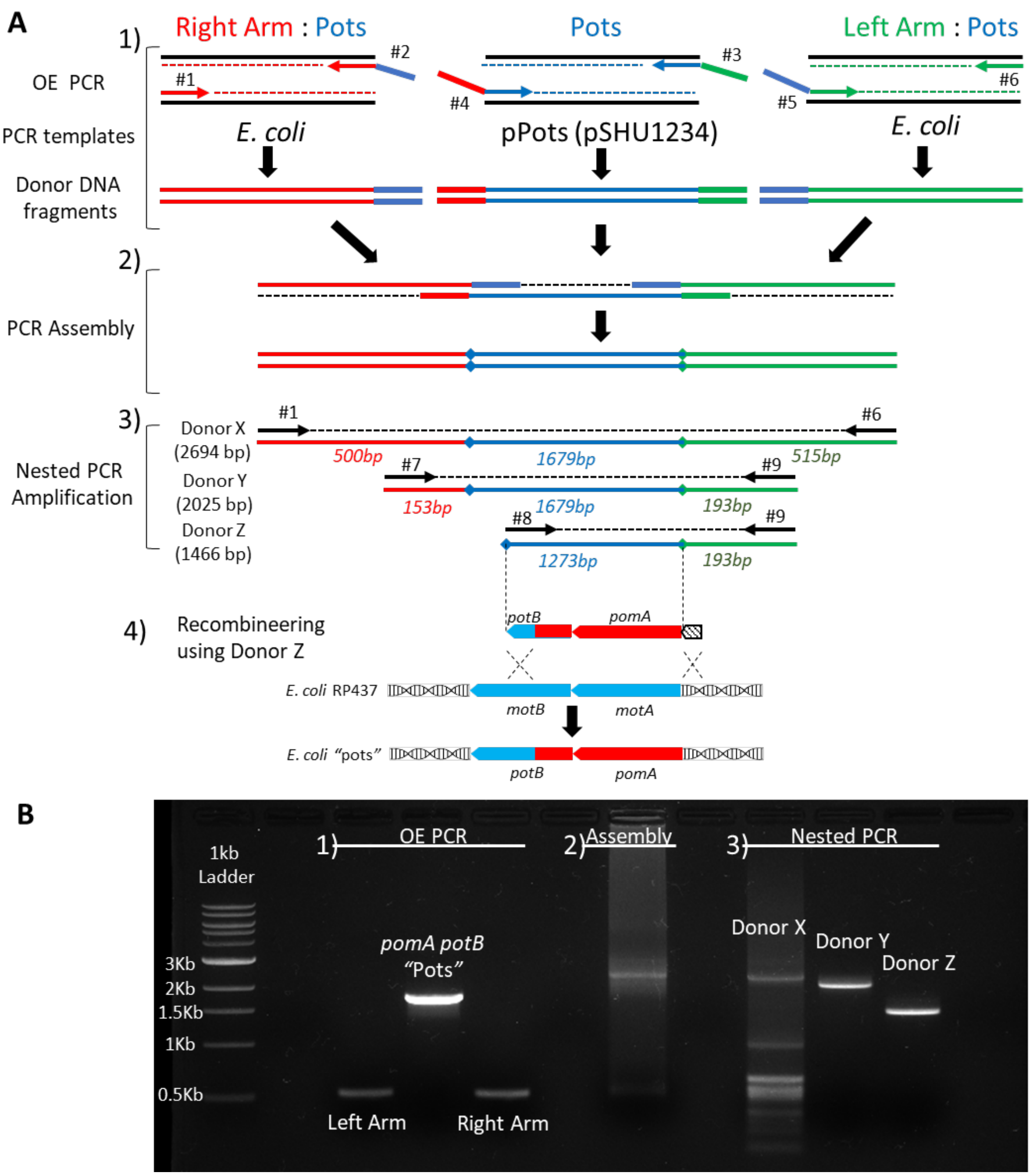


**Supplementary Fig. 2. Synthesis of double-stranded, linear donor DNA template.** A) Step-by-step graphical summary of the protocol employed to assemble the donor DNA for no-SCAR editing. 1: The overlap extension PCR technique (Higuchi, Krummel et al. 1988) was employed to assemble linear double stranded DNA molecules (dsDNA) using 3 starting fragments. These fragments included the up- and down-stream homology arms (right and left arm respectively, each 500 bp-long) intended to flank a third fragment encoding *pomApotB* (Pots) for *motAB* editing. Each dsDNA fragment was generated in a separate PCR reaction using primers bearing 5' overhangs to produce DNA that would be able hybridize with the appropriate flanking dsDNA fragment in the subsequent assembly step. Primers were designed to preserve the native gene orientation on the chromosome since *motAB* are encoded on the lagging strand. Primers are indicated by their ID# and sequences are provided in Supplementary Fig. 12. 2) equimolar amounts of each fragment were included in a PCR reaction mix without additional primers to assemble a continuous dsDNA molecule. 3) The resulting assembly was used as template for a nested PCR reaction to generate Donor DNA of different lengths to test in no-SCAR editing (full length Donor X: 2694 bp; Donor Y: 2025 bp; Donor Z: 1466 bp). The length of each fragment and the approximate position of the primers used in the nested PCR reaction is indicated in the diagram. 4) Diagram of the expected recombineering reaction using the shorter Donor Z fragment. The Right homology arm appears dispensable since the *motB* section of *potB* can act as the up-stream homology sequence during recombineering to generate a Pots-edited clone. B) Agarose gel electrophoresis of steps 1-3 described in A). The Donor DNA was gel extracted and purified before being electroporated into *E. coli* for editing.

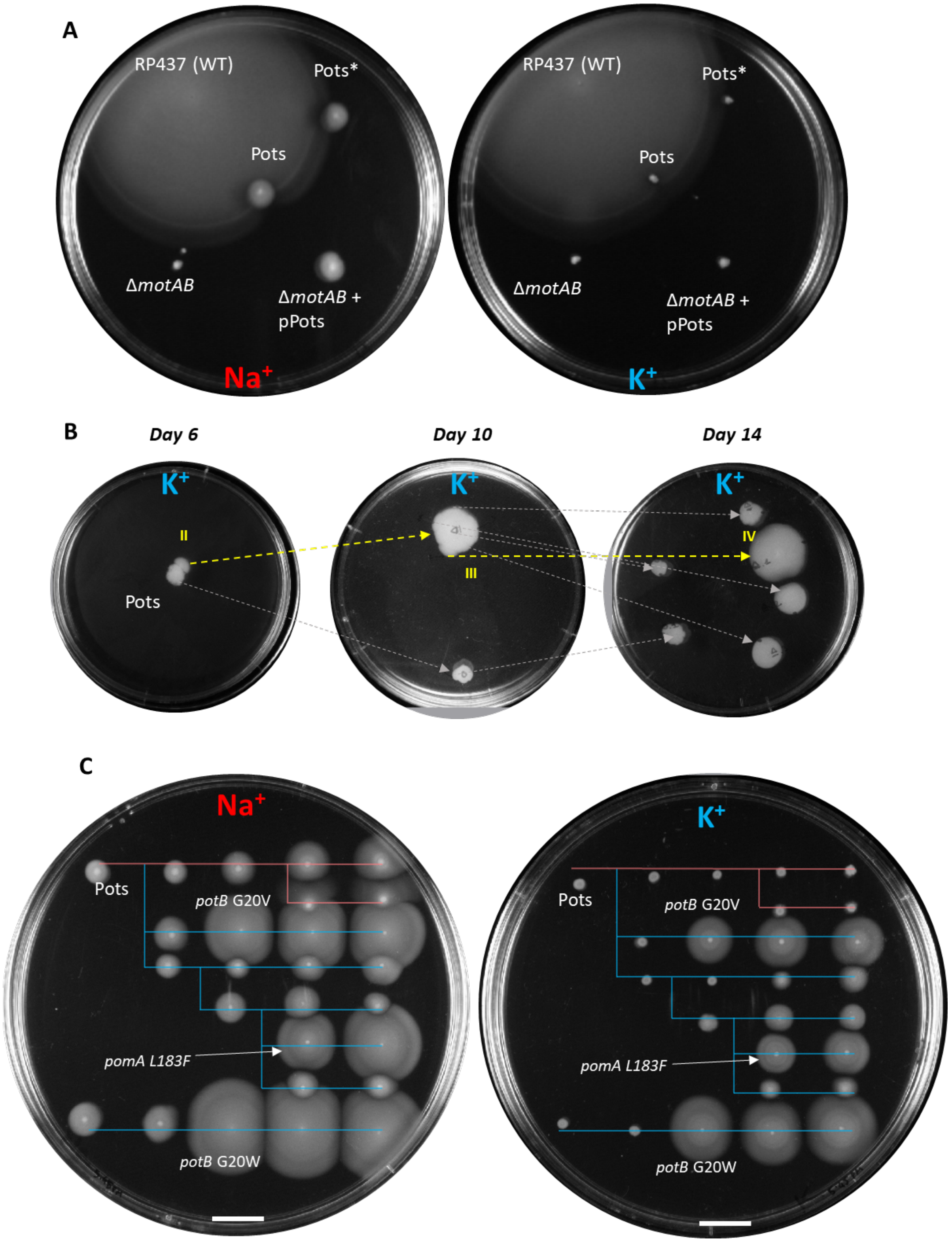


**Supplementary Fig. 3. Characterization and directed evolution of Sodium-powered *E. coli* Pots strain.** A) Soft agar swim plate assay to assess bacterial motility. The strains indicated above were inoculated on 0.3% Agar prepared with either $Na^+$LB (left) or $K^+$LB (right) media and incubated at 30°C for 24 hrs. The plates contain no antibiotics and 0.4% arabinose to induce expression from pPots. Strains: RP437 (WT, parent strain), Pots* (not-fully cured, carries plasmid pCas9cr4), Pots (cured of all plasmids), Δ*motAB* (E. coli RP6894) with or without pPots (pSHU1234, $Cm^+$, encoding the *pomApotB* construct under control of an inducible Arabinose promoter). B) Directed evolution experiment plates. A single Pots colony was inoculated on a $K^+$ soft agar plate and incubated at 30˚C until flares developed. The edge of the flare was then transferred onto a fresh plate and allowed to spread radially until at 4-day intervals. Yellow arrows indicate an improved swimmer subpopulation being propagated. Grey arrows indicate other portions of the swim ring being propagated. Flares labelled in yellow correspond to: II = L4.2, III = L5.3, IV = L6.4 in Fig. 2A, respectively. C) the recapitulation plates shown in Fig. 1C after a further 22 hr incubation at at 30°C.

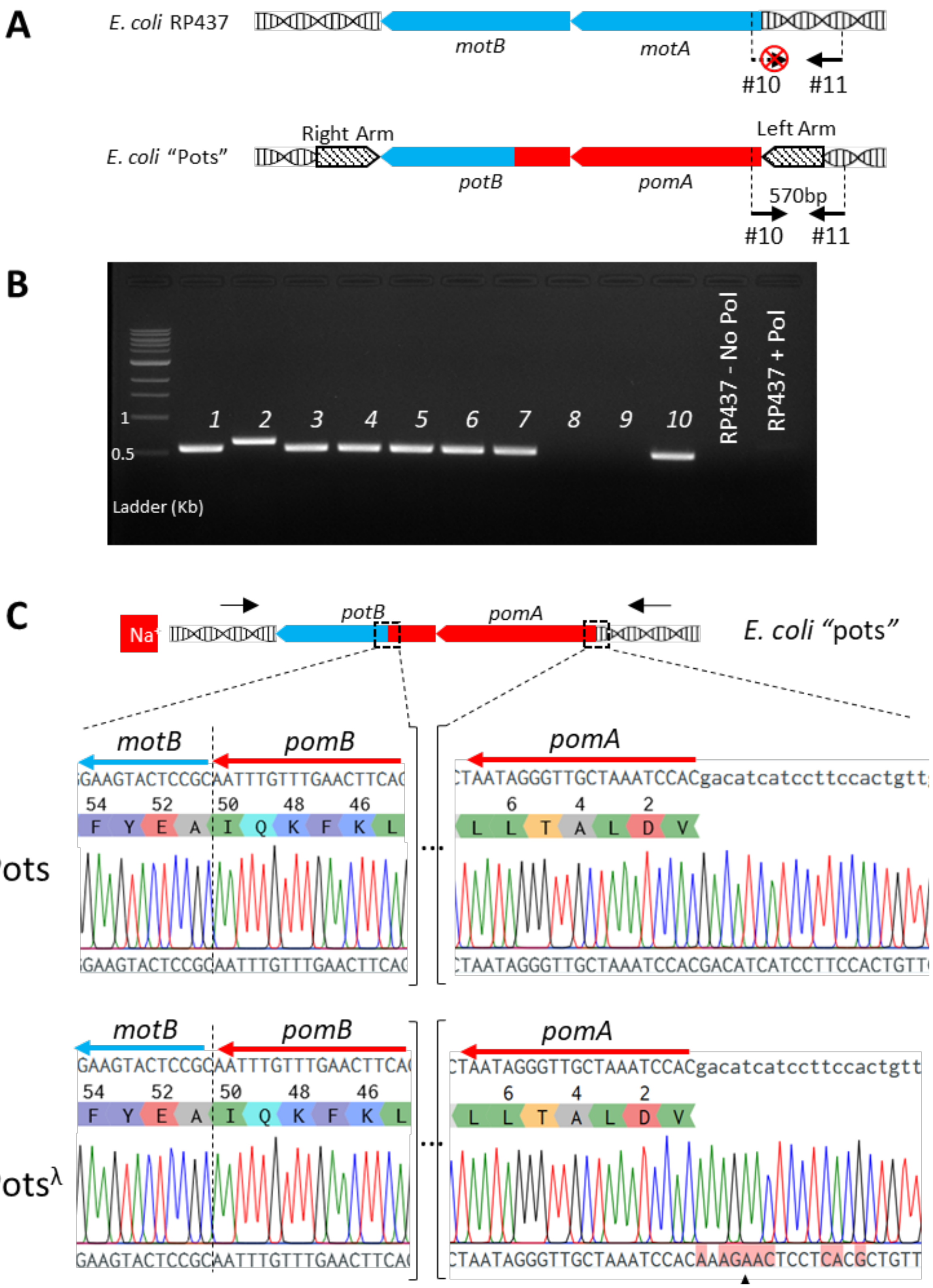


**Supplementary Fig. 4. Verification of successful genome editing using colony PCR and sequencing.** A) Cartoon diagram of the colony PCR reaction specific for Pots-edited clones. The primers indicated by the arrows are designed to amplify the section of chromosome spanning the 5' region of pomA and a section downstream of the Left homology arm to yield a 570bp amplicon. Primers are indicated by their ID#

and sequences are provided in Supplementary Fig. 10. B) agarose gel electrophoresis of 12x colony PCR reactions. Lanes 1-10 contain ampicons from candidate edited colonies, followed by the control reaction from WT RP437 with (+ Pol) and without (No-Pol) added Q5 polymerase. C) PCR products for Sanger sequencing were generated using primers flanking the Donor DNA recombination site. Chromatograms of sequenced Pots and Pots$^{\lambda}$ clones highlight the *motB/pomB* junction in potB and the 5' UTR region of *E. coli* RP437 motA. Pots retained the native *E. coli* sequence upstream of motA while Pots$^{\lambda}$ carried the *V. alginolyticus* sequence upstream of *pomA*. This clone was found to contain a single base mutation from C to A upstream from pomA at the site indicated by the black triangle.

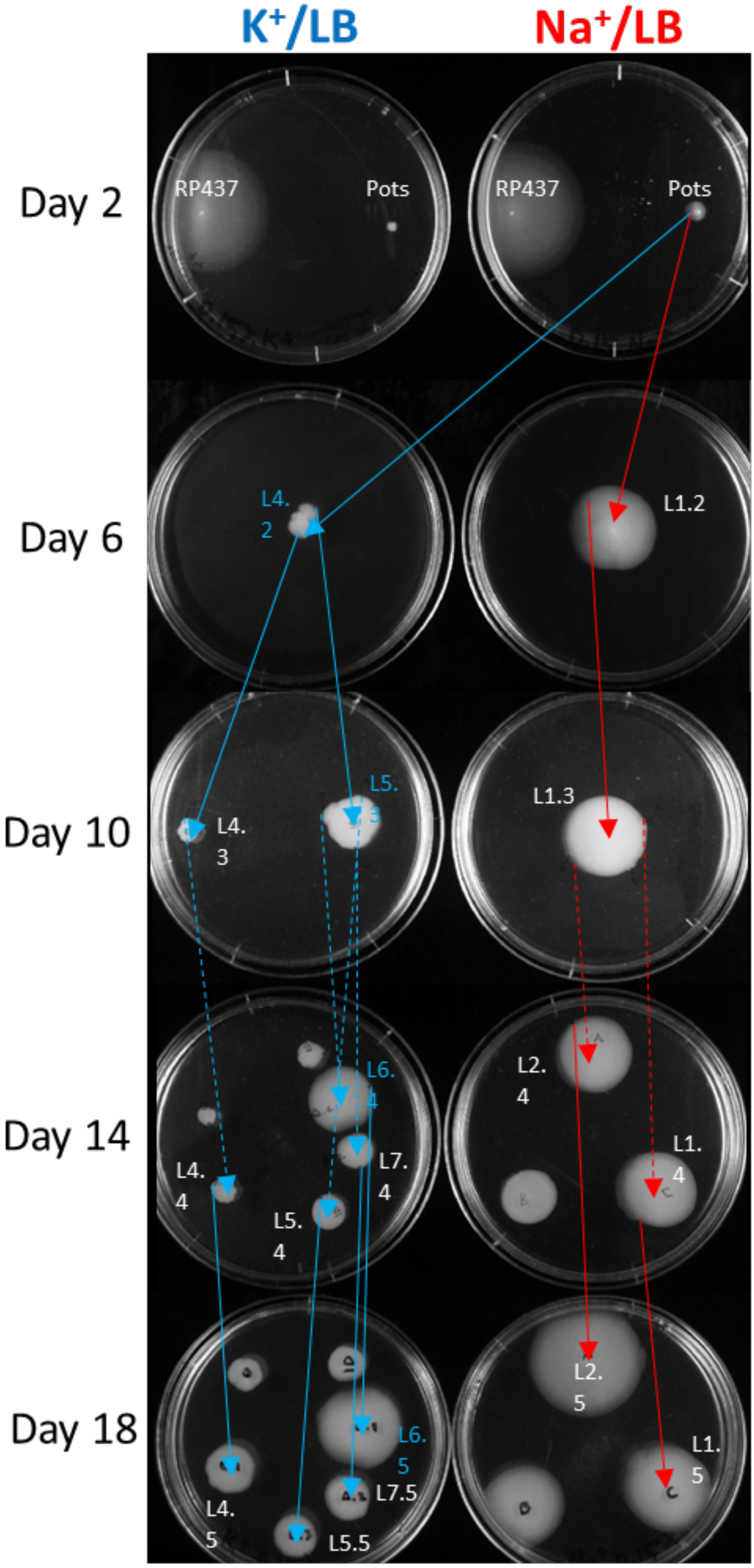


**Supplementary Fig. 5. Summary of the directed evolution experiment using Pots**. Each pair of $K^+$ and $Na^+$ swim plates was photographed immediately before passaging onto a fresh plate. Blue lines indicate passaging on $K^+$LB, red lines for $Na^+$LB. The colonies selected for whole genome sequencing (WGS) are labelled according to Fig. 2A. Colonies highlighted in cyan indicate members of the lineage which incorporated the *pomA* L183F mutation.

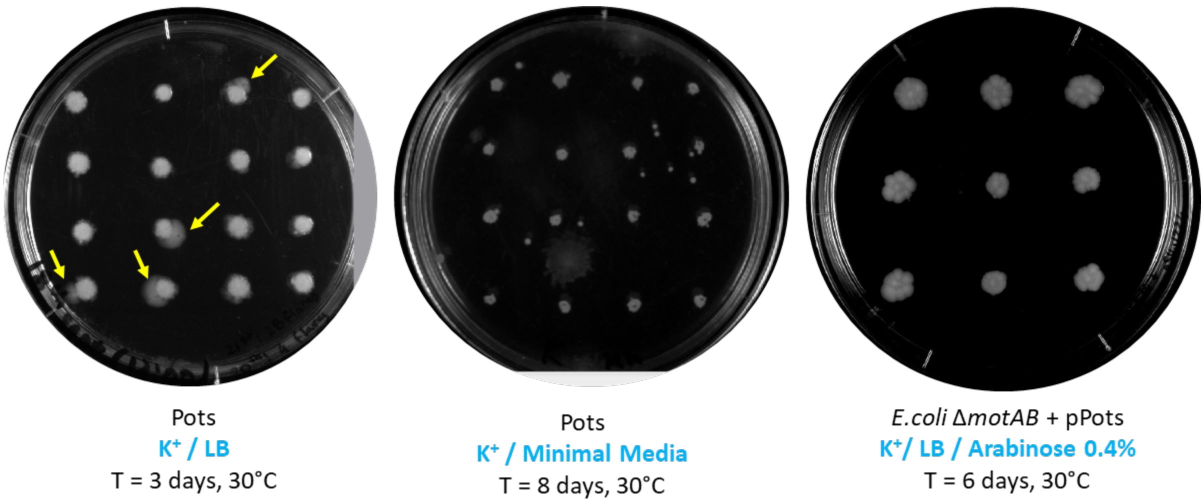


**Supplementary Fig. 6. Spontaneous flaring on different growth substrates.** Pots colonies were found to be able to produce flares on $K^+$/LB (yellow arrows) but not on $K^+$/ minimal media or when *pomA potB* was expressed from an Arabinose-inducible plasmid (pPots). The growth condition and length of incubation are indicated below each plate.

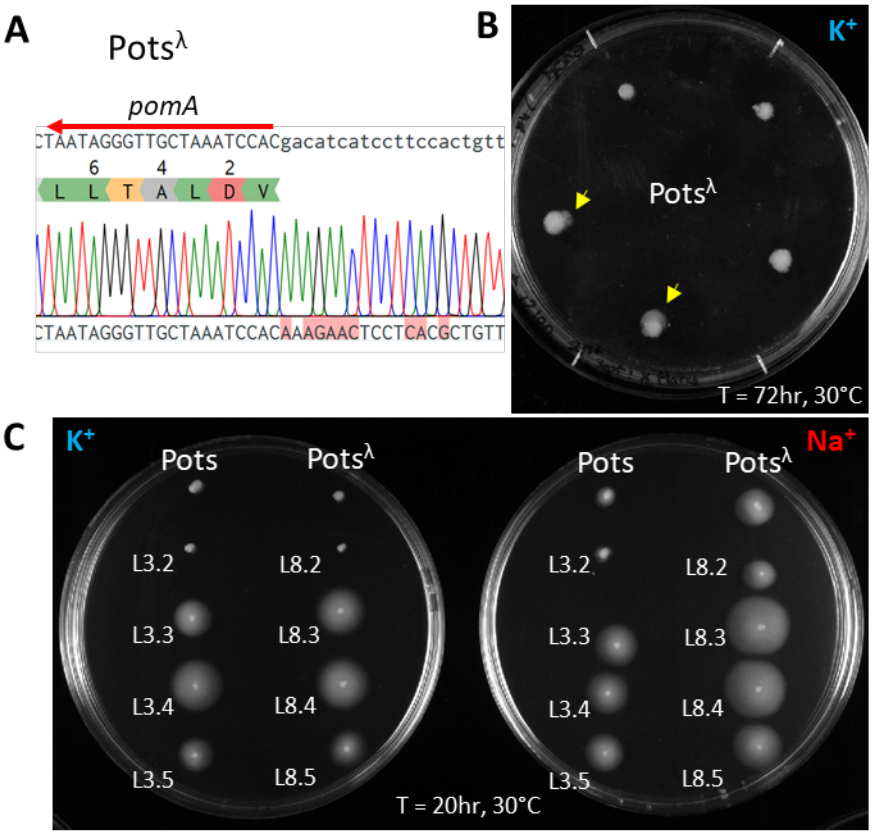


**Supplementary Fig. 7. Directed evolution of the Pots$^{\lambda}$ *E. coli* strain.** A) Pots$^{\lambda}$ sequencing results indicate the presence of the *pomA* 5' UTR region of *V. alginolyticus.* B) Pots$^{\lambda}$ colonies were able to produce flares (yellow arrows) on soft agar swim plate made of $K^+$LB within 3 days of incubation. C) Lineage recapitulation of a Pots and a Pots$^{\lambda}$ lineage which picked up mutations G20V (L3) and G20W (L8) in *potB,* respectively (see Fig. 2A).

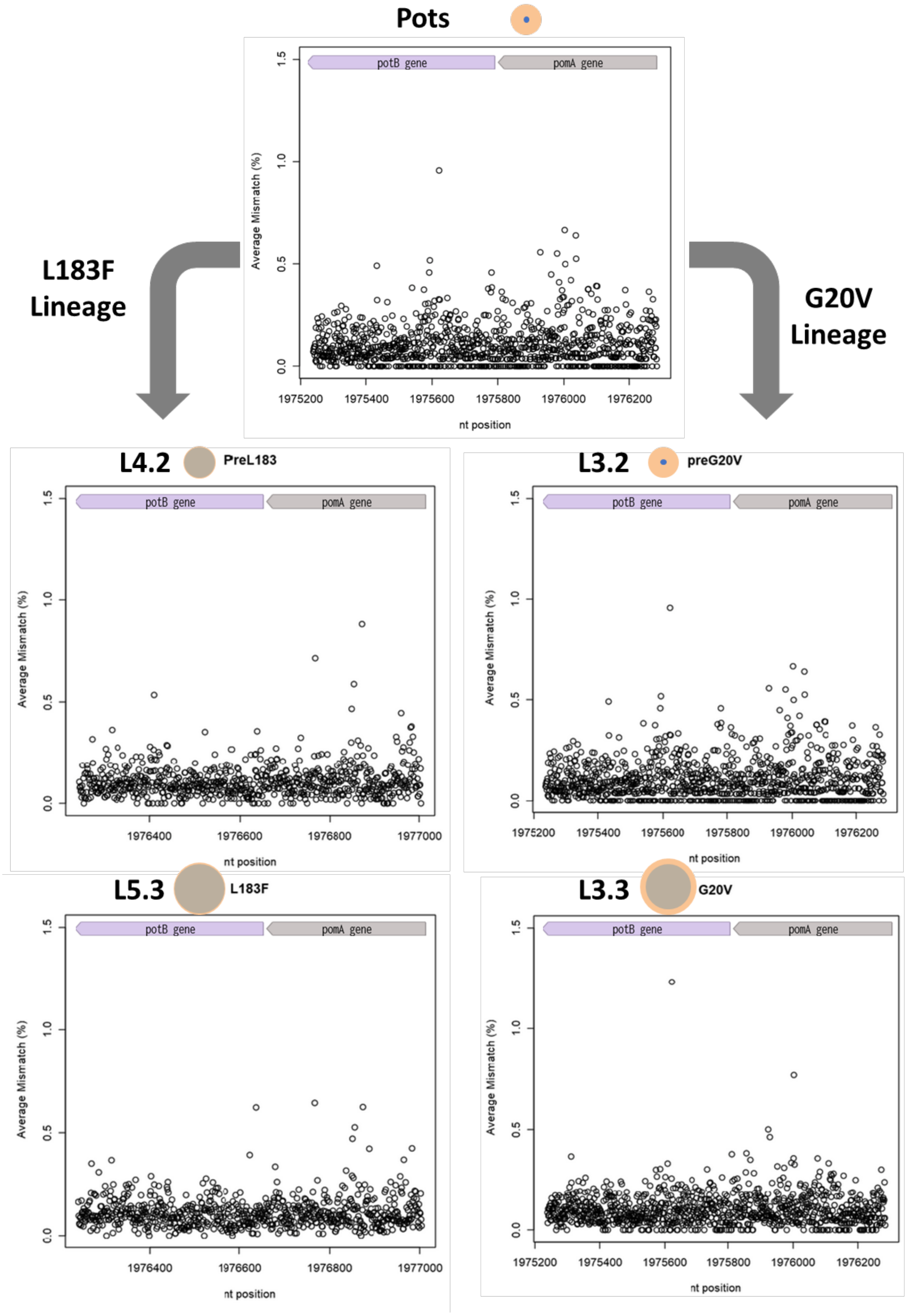


**Supplementary Fig. 8. Average mismatch percentage at each nucleotide position in *pomA* and *potB* RNA transcripts from Pots and evolved strains from the L183F and G20V lineages.** Each graph describes the nucleotide variation in all experimental replicates of the strain submitted for RNAseq. A cartoon of the gene orientation is displayed at the top of each graph. Concentric circles as presented in Fig.1B (orange: $Na^{+}$ motility; blue: $K^{+}$ motility) are shown next to each strain to indicted the motility phenotype displayed. Blue dots indicate lack of motility on $K^+$ swim agar. All nucleotide mismatches in mapped transcripts fall within the same range (0-1.5%). No enrichment of mutant transcript was found in strains L4.2 and L3.2, indicating that fixation of the mutation in the subsequent lineage member occurred independently of RNA modifications.

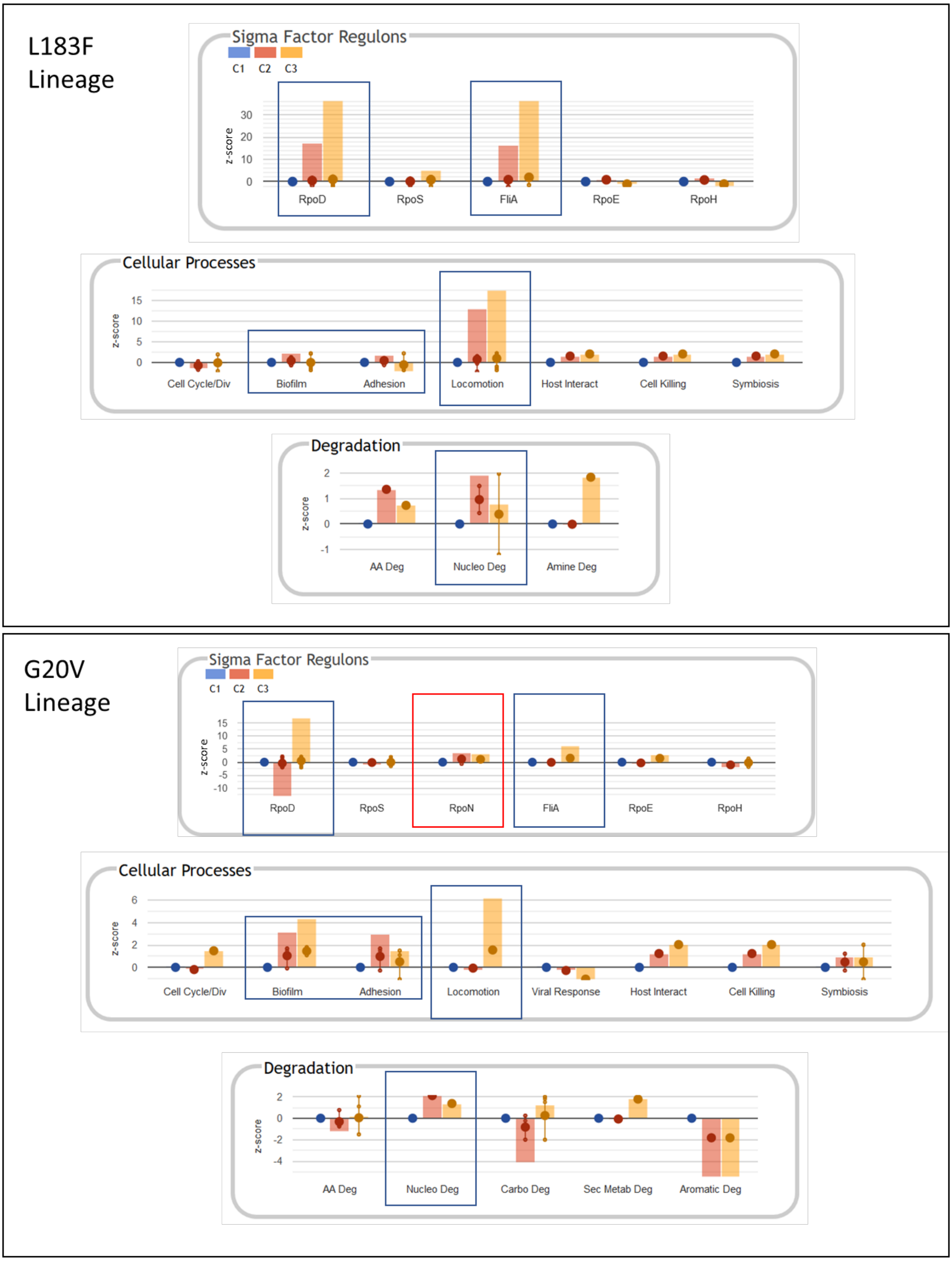


**Supplementary Fig. 9. Ecocyc analysis of RNAseq datasets.** Results from analysis of the L183F and G20V lineage datasets (top and bottom boxes, respectively) and presented as displayed in the Ecocyc Pathway Tools Omics Dashboard. These results were obtained by carrying out the analysis on the same z-score values as presented in heatmap format in fig.2A and 2B. Each box displays selected features presented in the Ecocyc analysis dashboard. Blue, red and yellow bars display activity levels calculated for Pots (C1, blue), the pre-mutation stage (C2, red) and the mutant stage of the lineage trio (C3, yellow). Top row: Predicted Sigma Factor regulon activity underlying the activities of the differentially expressed genes in each lineage in fig.2A and 2B. Coloured bars indicate the sum of all data values in the Sigma factor subsystem, which is indicative of the relative proportion of cell resources that are being invested in each subsystem. Different levels of sigma factor usage RpoD ($\sigma^{70}$), RpoN ($\sigma^{54}$) and FliA ($\sigma^{28}$) are highlighted by the blue and red boxes above each bar chart. Middle row: Predicted activity of various cellular processes of *E. coli*. Differences in the involvement of 'Biofilm', 'Adhesion' and 'Locomotion' subsystems are highlighted by boxes above the bar charts. Bottom row: Predicted activity of catabolic (Degradation) pathways specific to different substrates. Both lineages appear to upregulate Nucleotide degradation processes ('Nucleo Deg') at their pre-mutation stage (C2, red bar).

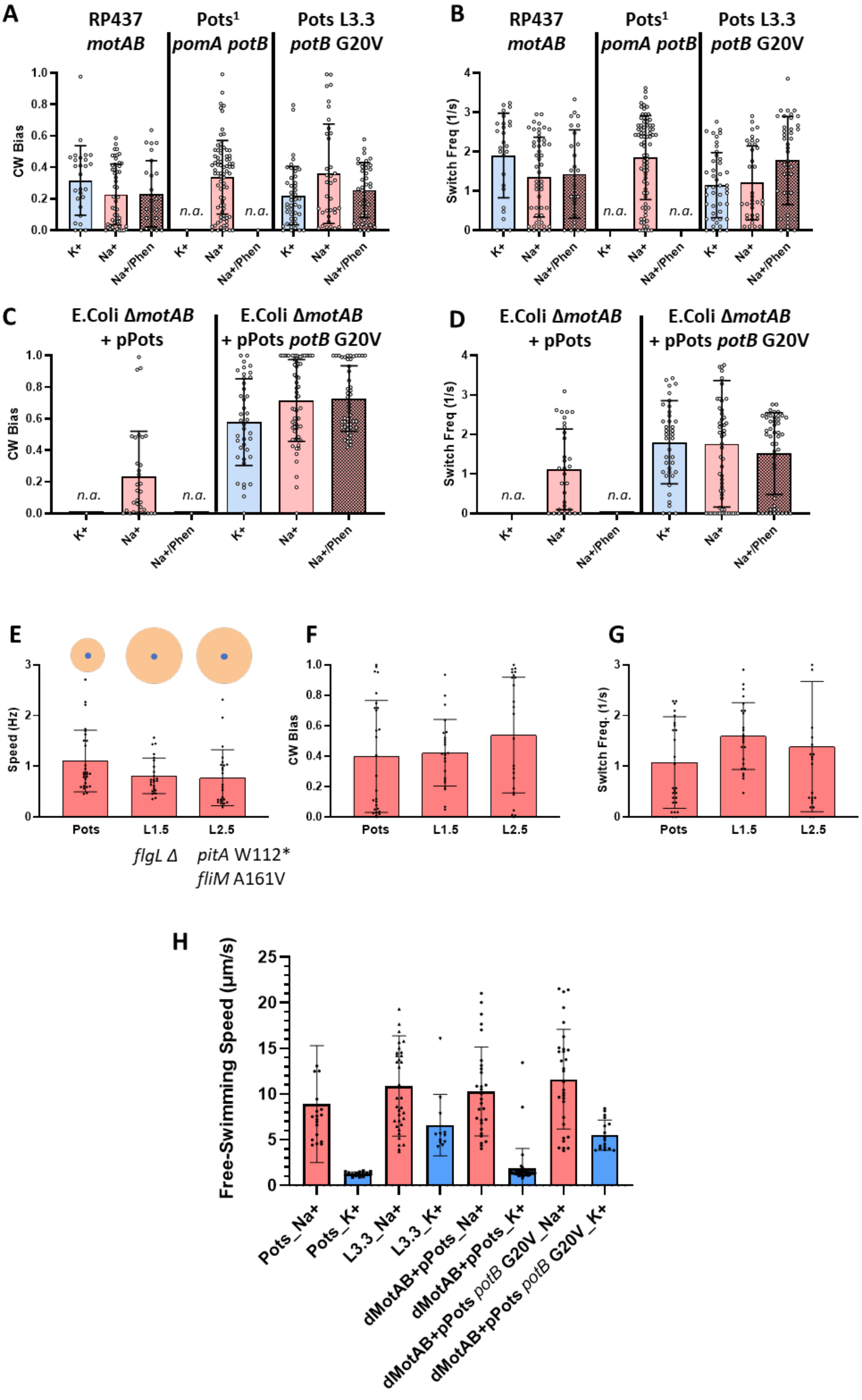
A
RP437
motAB
Pots1
pomA potB
Pots L3.3
potB G20V
CW Bias
K+
Na+
Na+/Phen
n.a.
B
Switch Freq (1/s)
C
E.Coli ΔmotAB
+ pPots
E.Coli ΔmotAB
+ pPots potB G20V
D
E
Speed (Hz)
Pots
L1.5
L2.5
flgL Δ
pitA W112*
fliM A161V
F
G
Switch Freq. (1/s)
H
Free-Swimming Speed (µm/s)
Pots_Na+
Pots_K+
L3.3_Na+
L3.3_K+
dMotAB+pPots_Na+
dMotAB+pPots_K+
dMotAB+pPots potB G20V_Na+
dMotAB+pPots potB G20V_K+

**Supplementary Fig. 10. Additional rotational parameters obtained from the tethered cell analysis software**. Blue bar indicates speed in 67 mM $K^+$ motility Buffer, red bar: 85 mM $Na^+$ motility Buffer; red patterned bar: 85 mM NaCl + 100 µM phenamil motility buffer. Clock-wise (CW) (A) bias and Switching Frequency (B) values for evolved strains described in Fig.3A. Number of cells analysed per condition (from left to right): RP437: 27, 51, 25; Pots: n.a, 78, n.a; Pots L3.3 *potB* G20V 45, 36, 39 (n.a. indicates no visible rotating cell). Error bars indicate S.D. Clockwise (CW) (C) bias and Switching Frequency (D) values for plasmid-transformed strains described in Fig.3B. Number of cells analysed per condition (from left to right): (Δ*motAB* + pPots: n.a., 32, n.a; Δ*motAB* + pPots *potB* G20V: 40, 63, 48). Error bars indicate SD. E-F-G) Tethered cell characterization of $Na^+$-adapted strains L1.5 and L2.5. Speed (E), CW Bias (F) and Switching frequency (G) were measured in 85 mM $Na^+$ motility buffer; A schematic representation of relative swim ring sizes and mutations associated with the L1.5 and L2.5 strains is also shown, extracted from Fig.1B. Number of cells analysed per condition: Pots (27), L1.5 (24), L2.5 (24). H) Mean free-swimming speed for untethered cells in $Na^+$ and $K^+$ motility buffers. Number of cells analysed per condition: Pots $Na^+$(19), Pots $K^+$ (16), L3.3 $Na^+$ (35), L3.3 $K^+$ (12), Δ*motAB* + pPots $Na^+$ (29), Δ*motAB* + pPots $K^+$ (42), Δ*motAB* + pPots *potB* G20V $Na^+$ (31), Δ*motAB* + pPots *potB* G20V $K^+$ (17). Error bars indicate Standard Deviation.

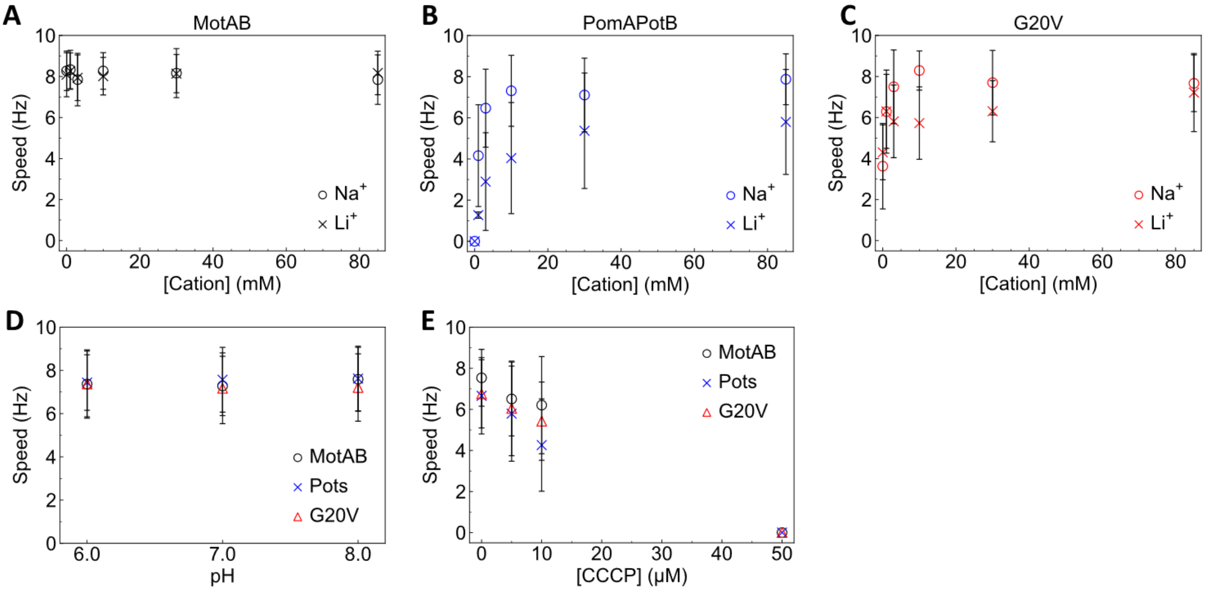


**Supplementary Fig. 11. Energisation curves, effect of pH and CCCP for MotA/MotB, PomA/PotB and PomA/PotB G20V stators.** A-C) Averaged tethered cell rotational speed in varying sodium (circle) and lithium (cross) concentration (energisation curves) for motors bearing the MotAB (A), PomA/PotB (Pots) (B) and PomA/PotB G20V (C) stators in presence of mixed cations according to the following equation [KCl] + ([NaCl] or [LiCl]) = 85 mM. The dependency of each motor on $Na^+$ or $Li^+$ was tested at 0, 1, 3, 10, 30 and 85 mM of ion respectively. The cations [85 mM] were mixed in motility buffer containing 10 mM KPi, 0.1 mM EDTA-2K, pH = 7.0. Each data point represents the average speed from a minimum of 25 cells (max N = 57 cells). Error bars indicate Standard Deviation (S.D.). D) Effect of motility buffer (10 mM KPi, 0.1 mM EDTA-2K, 85 mM NaCl) adjusted to pH 6, 7 or 8 on motor speed. Each data point represents the average speed from a minimum of 39 cells (max N = 98 cells) . E) Effect of protonophore CCCP on motor speed. CCCP was tested in motility buffer (10 mM KPi, 0.1 mM EDTA-2K, 85 mM NaCl, pH = 7.0) at 0, 5, 10 and 50 μM. Each data point represents the average speed from a minimum of 41 cells (max N = 95 cells). All tethered cell assays were performed on FliC-sticky cells expressing the stator sets from a pPots plasmid.

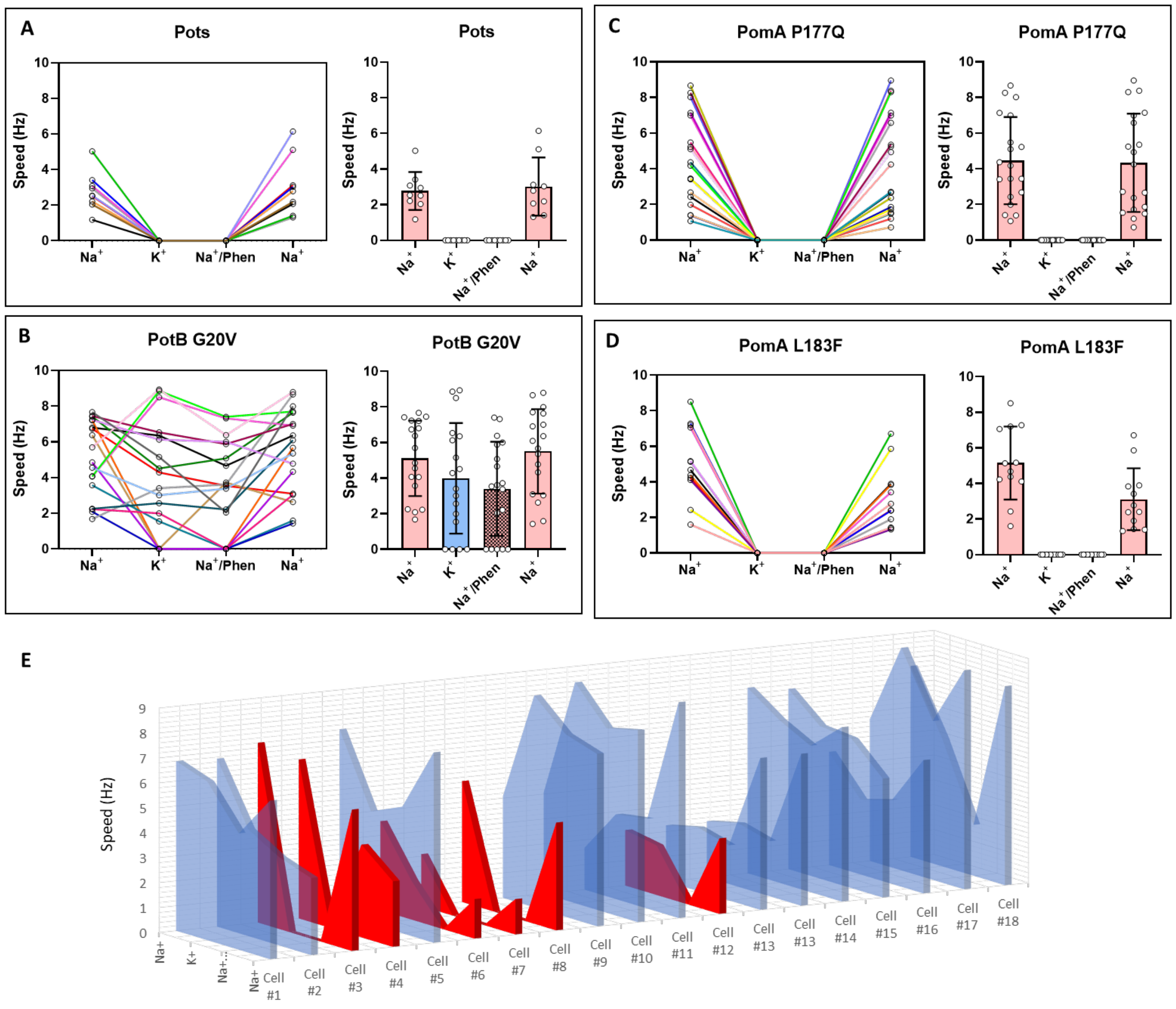


**Supplementary Fig. 12. Tracking single tethered cells during motility buffer exchange.** The rotational speed of lineage members (A) Pots, (B) *potB* G20V (L3.3), (C) *pomA* P177Q and (D) *pomA* L183F (L6.4) was analysed in detail using the tethered cell assay while sequentially exchanging motility buffers and tracking the behaviour of single tethered cells. Each box displays the single cell speed measured under each buffer condition (left). Coloured lines are used to track individual cells. The tethered cells were sequentially exposed to 85 mM $Na^+$, 85 mM $K^+$, 85 mM NaCl + 100 µM phenamil and 85 mM $Na^+$ motility buffers, in this order. Only cells which were motile after the last $Na^+$ buffer exchange were included in the analysis. The average speed value of all analysed cells per condition is quantified in the bar chart on the right hand-side of each box (error bars represent SD). The red bar indicates speed in 85 mM $Na^+$ motility buffer, blue bar: 85 mM $K^+$ motility buffer; red patterned bar: 85 mM NaCl + 100 µM phenamil motility buffer. Number of cells analysed per strain (from A to D): Pots (9), *potB* G20V (18), *pomA* P177Q (19) and *pomA* L183F (12). The tethered cell assay videos were acquired at 60 frame/s. E) 3D plot of cell rotation speed values for each cell plotted in panel B. Cells that stopped rotating when $K^+$MB or $Na^+$MB + phenamil was tested are highlighted in red. Cells that maintained motility under all motility buffer conditions are highlighted in blue.

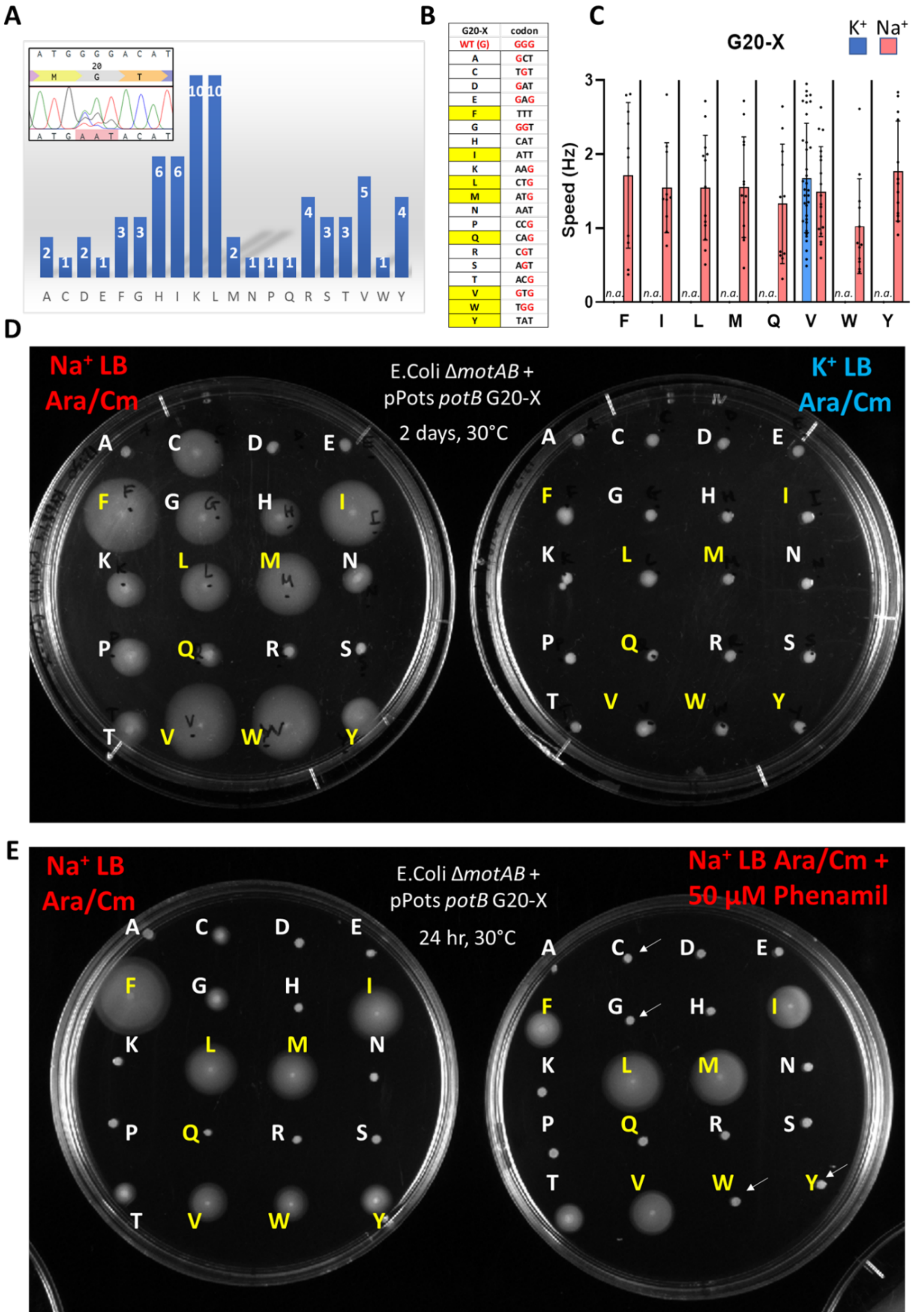


**Supplementary Fig. 13. Saturation Mutagenesis of the PotB G20 site.**

A) The ‘22-c trick’ technique was employed to generate all possible amino acid substitutions at the PotB G20 site on the pPots plasmid in a single step. The bar chart indicates the frequency of each AA substitution across 70 screened colonies after transformation with a randomly mutagenized plasmid population (Sanger chromatogram shown in inset highlights the nucleotide degeneracy at the G20 codon of PotB). B) list of selected clones for each AA substitution and their respective codon. The WT G20 and its respective codon (GGG) are indicated at the top of the list. Codon letters highlighted in red indicate unchanged nucleotides after mutagenesis. The AA substitutions highlighted in yellow were selected for functional characterization based on their ability to swim on soft agar (panel D). C) Tethered cell assay of Δ*motAB E.*

*coli* strain trasnsformed with pPots *potB* G20-F, I, L, M, Q, V, W and Y. Average tethered cell rotation speed in $K^+$ (blue bar) or $Na^+$ (red) motility buffer (MB). Only G20V was found to rotate in K+ motility buffer. Number of cells analysed per condition: F $^{Na+}$ 10, I $^{Na+}$ 1, L $^{Na+}$ 13, M $^{Na+}$ 13, Q $^{Na+}$ 11, V $^{K+}$ 34, V $^{Na+}$ 18, W $^{Na+}$ 12, Y $^{Na+}$ 13 (n.a. indicates no visible rotating cell). Error bars indicate Standard Deviation (S.D.) D) Soft agar swim plates (ø = 10 cm) inoculated with *E. Coli* Δ*motAB* cells carrying the pPots *potB* G20-X mutants indicated in Panel B. Single colonies from a transformation plate were tapped onto $Na^+$ (red, left) or $K^+$ (blue, right) swim agar supplied with Arabinose and Chloramphenicol (Ara/Cm) for induction and selection, and incubated at 30°C for 48 hrs. E) Effect of 50 μM phenamil on *potB* G20-X mutants. White arrows point at G20 variants which appear sensitive to phenamil blockage (WT, G20C, G20W and G20Y). Soft agar swim plates were inoculated with the same strains as panel D and incubated for 24hr.

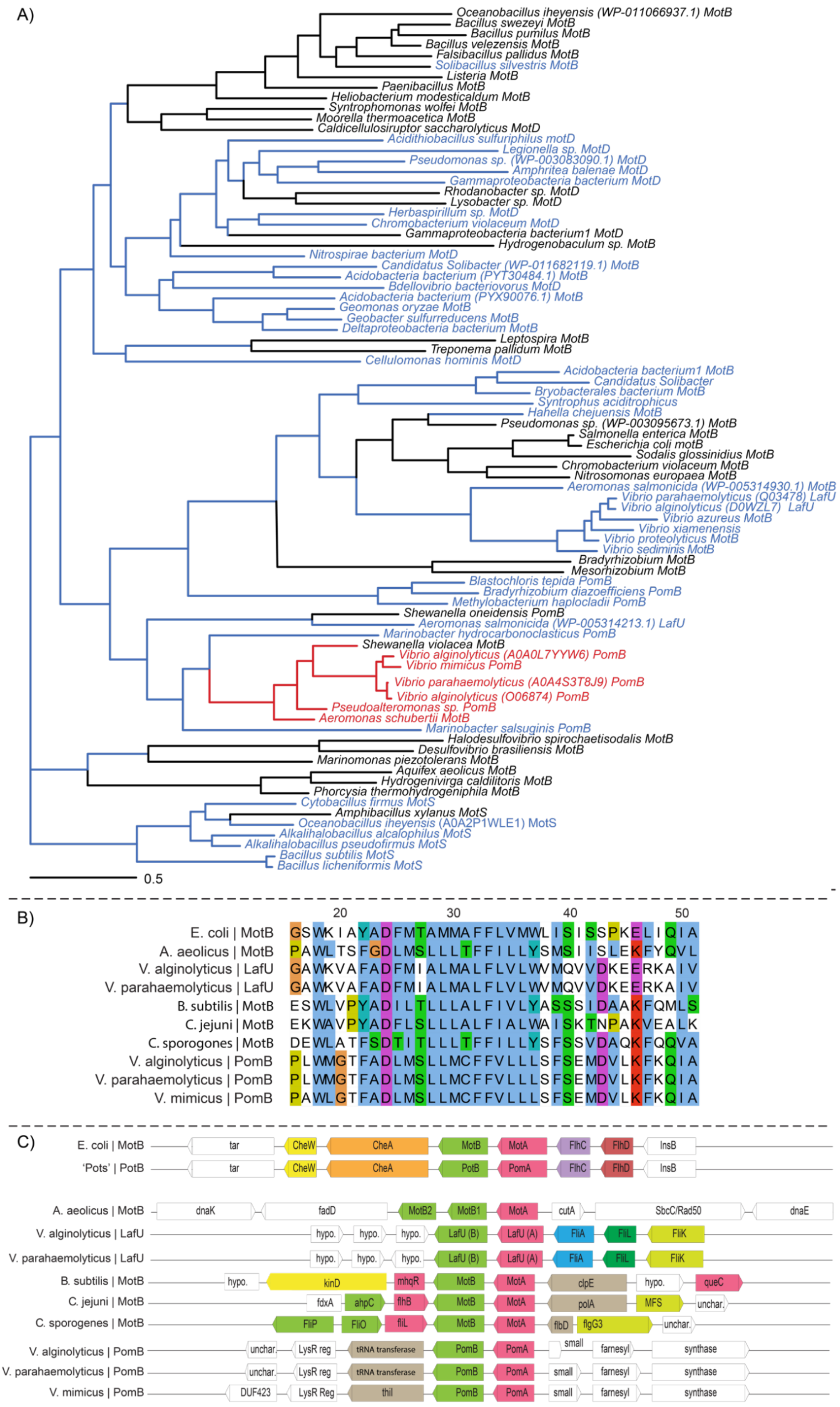

A)
Oceanobacillus iheyensis (WP-011066937.1) MotB
Bacillus swezeyi MotB
Bacillus pumilus MotB
Bacillus velezensis MotB
Falsibacillus pallidus MotB
Solibacillus silvestris MotB
Listeria MotB
Paenibacillus MotB
Heliobacterium modesticaldum MotB
Syntrophomonas wolfei MotB
Moorella thermoacetica MotB
Caldicellulosiruptor saccharolyticus MotD
Acidithiobacillus sulfuriphilus motD
Legionella sp. MotD
Pseudomonas sp. (WP-003083090.1) MotD
Amphritea balenae MotD
Gammaproteobacteria bacterium MotD
Rhodanobacter sp. MotD
Lysobacter sp. MotD
Herbaspirillum sp. MotD
Chromobacterium violaceum MotD
Gammaproteobacteria bacterium1 MotD
Hydrogenobaculum sp. MotB
Nitrospirae bacterium MotD
Candidatus Solibacter (WP-011682119.1) MotB
Acidobacteria bacterium (PYT30484.1) MotB
Bdellovibrio bacteriovorus MotD
Acidobacteria bacterium (PYX90076.1) MotB
Geomonas oryzae MotB
Geobacter sulfurreducens MotB
Deltaproteobacteria bacterium MotB
Leptospira MotB
Treponema pallidum MotB
Cellulomonas hominis MotD
Acidobacteria bacterium1 MotB
Candidatus Solibacter
Bryobacterales bacterium MotB
Syntrophus aciditrophicus
Hahella chejuensis MotB
Pseudomonas sp. (WP-003095673.1) MotB
Salmonella enterica MotB
Escherichia coli motB
Sodalis glossinidius MotB
Chromobacterium violaceum MotB
Nitrosomonas europaea MotB
Aeromonas salmonicida (WP-005314930.1) MotB
Vibrio parahaemolyticus (Q03478) LafU
Vibrio alginolyticus (D0WZL7) LafU
Vibrio azureus MotB
Vibrio xiamenensis
Vibrio proteolyticus MotB
Vibrio sediminis MotB
Bradyrhizobium MotB
Mesorhizobium MotB
Blastochloris tepida PomB
Bradyrhizobium diazoefficiens PomB
Methylobacterium haplocladii PomB
Shewanella oneidensis PomB
Aeromonas salmonicida (WP-005314213.1) LafU
Marinobacter hydrocarbonoclasticus PomB
Shewanella violacea MotB
Vibrio alginolyticus (A0A0L7YYW6) PomB
Vibrio mimicus PomB
Vibrio parahaemolyticus (A0A4S3T8J9) PomB
Vibrio alginolyticus (O06874) PomB
Pseudoalteromonas sp. PomB
Aeromonas schubertii MotB
Marinobacter salsuginis PomB
Halodesulfovibrio spirochaetisodalis MotB
Desulfovibrio brasiliensis MotB
Marinomonas piezotolerans MotB
Aquifex aeolicus MotB
Hydrogenivirga caldilitoris MotB
Phorcysia thermohydrogeniphila MotB
Cytobacillus firmus MotS
Amphibacillus xylanus MotS
Oceanobacillus iheyensis (A0A2P1WLE1) MotS
Alkalihalobacillus alcalophilus MotS
Alkalihalobacillus pseudofirmus MotS
Bacillus subtilis MotS
Bacillus licheniformis MotS
0.5
B)
20
30
40
50
E. coli | MotB GSWKIAYADFMTAMMAFFLVMWLISISSPKELIQIA
A. aeolicus | MotB PAWLTSFGDLMSLLLTFFILLYSMSIISLEKFYQVL
V. alginolyticus | LafU GAWKVAFADFMIALMALFLVLWMQVVDKEERKAIV
V. parahaemolyticus | LafU GAWKVAFADFMIALMALFLVLWMQVVDKEERKAIV
B. subtilis | MotB ESWLVPYADILTLLLALFIVLYASSSIDAAKFQMLS
C. jejuni | MotB EKWAVPYADFLSLLLALFIALWAISKTNPAKVEALK
C. sporogones | MotB DEWLATFSDTITLLLTFFILLYSFSSVDAQKFQQVA
V. alginolyticus | PomB PLWMGTFADLMSLLMCFFVLLLSFSEMDVLKFKQIA
V. parahaemolyticus | PomB PLWMGTFADLMSLLMCFFVLLLSFSEMDVLKFKQIA
V. mimicus | PomB PAWLGTFADLMSLLMCFFVLLLSFSEMDVLKFKQIA
C)
E. coli | MotB
tar CheW CheA MotB MotA FlhC FlhD InsB
'Pots' | PotB
tar CheW CheA PotB PomA FlhC FlhD InsB
A. aeolicus | MotB
dnaK fadD MotB2 MotB1 MotA cutA SbcC/Rad50 dnaE
V. alginolyticus | LafU
hypo. hypo. hypo. LafU (B) LafU (A) FliA FliL FliK
V. parahaemolyticus | LafU
hypo. hypo. hypo. LafU (B) LafU (A) FliA FliL FliK
B. subtilis | MotB
hypo. kinD mhqR MotB MotA clpE hypo. queC
C. jejuni | MotB
fdxA ahpC flhB MotB MotA polA MFS unchar.
C. sporogenes | MotB
FliP FliO fliL MotB MotA flbD flgG3 unchar.
V. alginolyticus | PomB
unchar. LysR reg tRNA transferase PomB PomA small farnesyl synthase
V. parahaemolyticus | PomB
unchar. LysR reg tRNA transferase PomB PomA small farnesyl synthase
V. mimicus | PomB
DUF423 LysR Reg thiI PomB PomA small farnesyl synthase

**Supplementary Fig. 14. Phylogenetic context of G20V mutation.** A) Phylogeny of 82 representative MotB/MotD/MotS/LafU/PomB proteins including select sodium and proton swimmers. Glycine/Valine identity at residue is indicated by text colour (blue: V20, pink: G20, black: other). Estimated sequence identity at each node indicated by branch colour (blue: V20, pink: G20, black: other, where two types meet the predicted residue at that node is indicated by the left branch). B) Sequence context around the transmembrane domain of the MotB subunit for *E. coli, A. aeolicus, B. subtilis, C. jejuni* and *C. sporogenes* MotB, PomB from *V. alginolyticus, V. parahaemolyticus* and *V. mimicus*, as well as LafU from *V. alginolyticus* and *V. parahaemolyticus*. In the *Vibrio* sp., sodium powered PomB is G20 whereas proton powered LafU is V20, showing a split at the critical residue adaptation characterised in this work. Colouring uses ClustalX scheme. For a comparable alignment of MotA species please consult Fig 3 in Biquet-Bisquert et al (Biquet-Bisquert et al., 2021). C) Genomic context around the PomA/PomB genes and MotA/MotB (LafU) genes in the species in B. In *E. coli* and in our edited strain 'Pots', *motA* is in proximity to regulatory genes *flhC* (controlled by master regulator *lrhA*), however in other strains such as the sodium powered *Vibrio* sp., the genomic context is different, with no signature in the genome that can be used to easily discern preference for proton- or sodium- power. Legend for non-flagellar genes in (B) in order of appearance (top to bottom, left to right): *tar:* methyl-accepting chemotaxis protein II, aspartate sensor receptor*, InsB:* insertion element IS1 protein InsB*, dnaK:* molecular chaperone DnaK*, fadD:* long-chain acyl-CoA synthetase*, cutA:* periplasmic divalent cation tolerance protein*, SbcC/Rad50:* (hypothetical) DNA repair protein SbcC/Rad50*, dnaE:* DNA polymerase III subunit alpha*, kinD:* sporulation kinase D, *mhqR*: MarR family transcriptional regulator, *ClpE:* ATP-dependent Clp protease ATP-binding subunit ClpE, *queC:* 7-cyano-7-deazaguanine synthase QueC, *fdxA:* ferredoxin, *ahpC:* alkyl hydroperoxide reductase, *polA*: DNA polymerase I, *MFS:* MFS transport protein, *LysR reg:* LysR family transcriptional regulator, glycine cleavage system transcriptional activator*, tRNA transferase:* tRNA uracil 4-sulfurtransferase, *small:* exodeoxyribonuclease VII small subunit*, farnesyl:* farnesyl diphosphate synthase*, synthase:* 1-deoxy-D-xylulose-5-phosphate synthase*, DUF423:* DUF423 domain-containing protein*, thiI:* tRNA uracil 4-sulfurtransferase. Hypo. Indicates hypothetical protein; unchar. indicates uncharacterised protein. Genomic context taken from KEGG database as per methods.

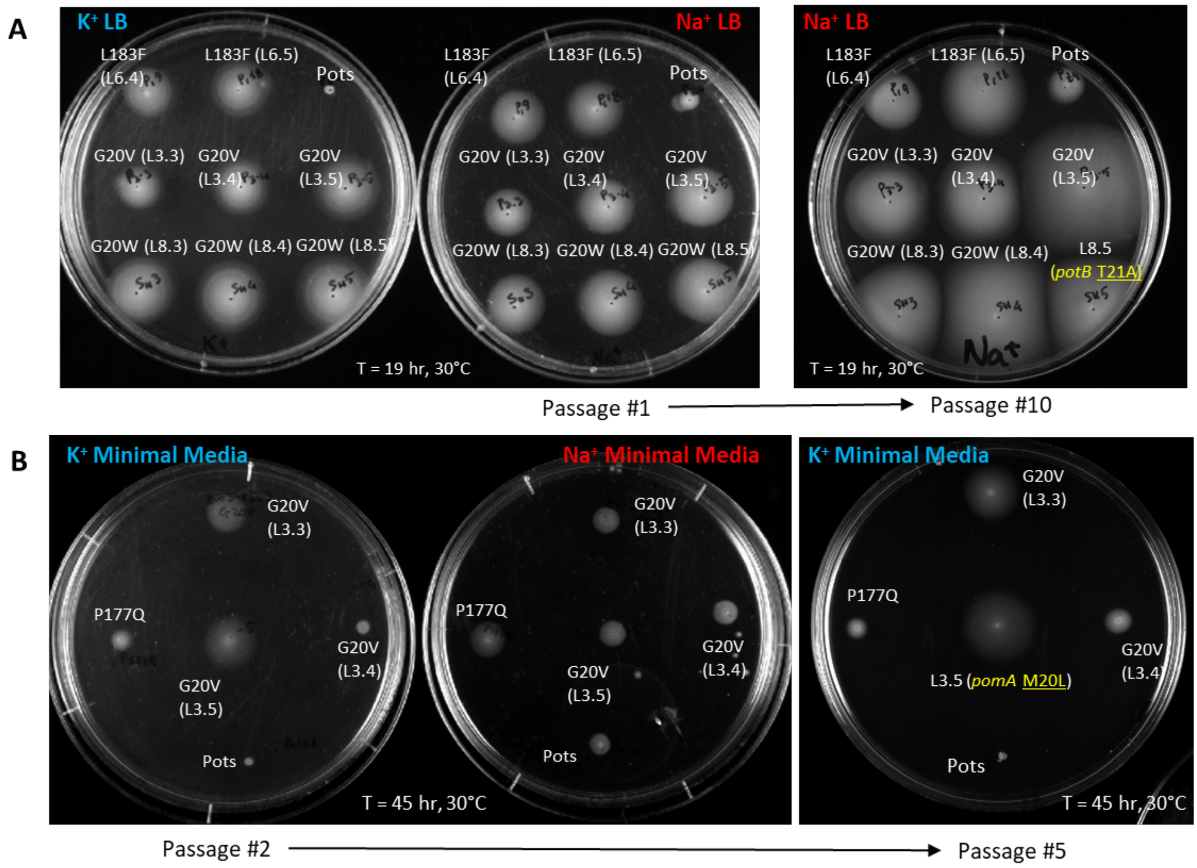


**Supplementary Fig. 15. Reversion testing and additional rounds of minimal media directed evolution.** A) The strains previously selected for WGS were tested for their ability to restore their mutant site to the original wild-type amino acid after being reintroduced to a sodium-rich environment. The mutant lineage members of Fig. 1B are shown here on $K^+$ and $Na^+$LB for comparison (passage #1). The edge of each swim ring was passaged daily on $Na^+$LB soft agar for a total of 10 passages and the final passage is shown on the right. Only the descendant of L8.5 was found to have incorporated a new mutation in *potB* (T21A). B) Stator protein mutants obtained during directed evolution on $K^+$LB were introduced to $K^+$ Minimal Media soft agar and passaged at 3-4 day intervals as in previous experiments. The strains which produced flares during the first incubation (*pomA* P177Q and *potB* G20V strains) were passaged and are shown here on $K^+$ and $Na^+$ Minimal Media for comparison (passage #2). After a total of 5 passages on $K^+$ Minimal Media, the descendant of L3.5 was found to have incorporated a new mutation in *pomA* (M20L).

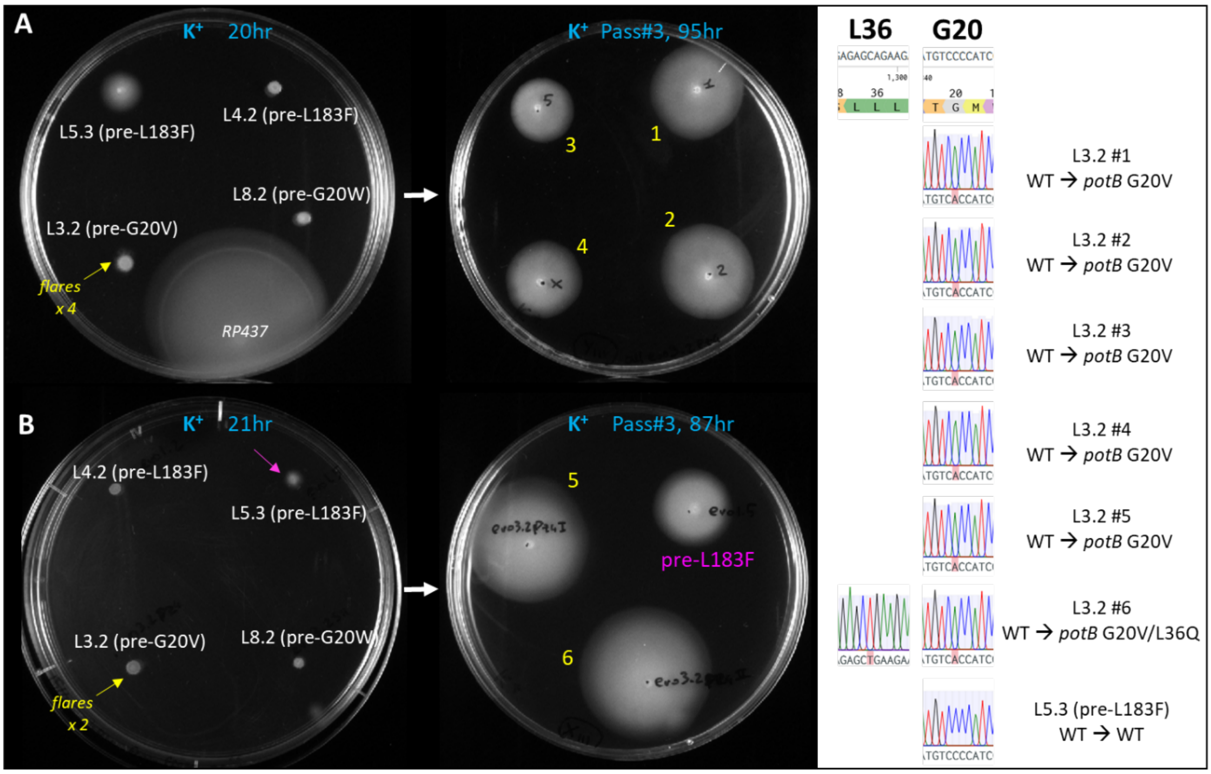


**Supplementary Fig. 16. Re-inoculation of L3.2 (pre-G20V) on K$^+$ swim agar produces flares and yields G20V mutation.** Frozen stocks (1 µl) of strains preceding the fixation of a stator mutant (L4.2 and L5.3 pre-*pomA* L183F, L3.2 pre-*potB* G20V and L8.2 pre-*potB* G20W) were re-inoculated on K$^+$LB swim plates and allowed to flare. Clones L5.3 and L3.2 were found to produce flares within 24hr in two separate experiments (A and B) for a total of 6x L3.2 descendants (yellow arrows, #1-6) and a single L5.3 descendant (pink arrow, pre-L183F). These flares were passaged twice more for the subsequent 2 days and screened for stator mutations by Sanger sequencing colony PCR products. The Chromatograms from sequencing reactions are shown for nucleotides encompassing the L36 and G20 sites of potB. All 6 pre-*potB* G20V were found to have developed the same *potB* G20V mutation (ccc → cac), and clone #6 also displayed the *potB* L36Q mutation, also previously found (Fig.3C). In contrast, the single L5.3 descendant did not display any mutation in the stator genes *pomA* and *potB*.

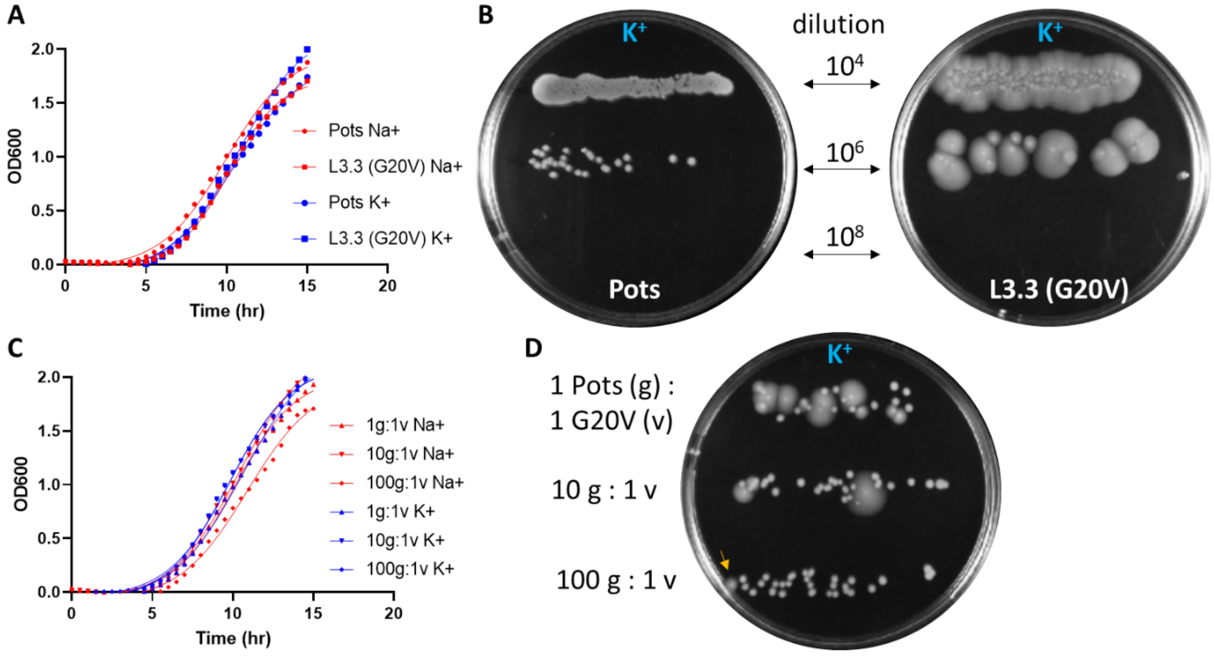


**Supplementary Fig. 17. Growth parameters and fitness comparison between Pots and L3.3 (G20V) strains.** A) Optical density ($OD_{600}$) determination of growth curves of Pots and L3.3 strains growing at 30°C in either $Na^+$- or $K^+$-based LB broths. B) Pots an and L3.3 cultures grown in $K^+$LB (OD600=0.25) diluted $10^4$-fold, $10^6$-fold, $10^8$-fold and inoculated on $K^+$LB swim plates (20 µl streaks) for 48hr at 30°C. Comparison of $10^6$ dilutions shows how all Pots colonies are non-motile while all L3.3 colonies display swim rings. C) Optical density ($OD_{600}$) determination of growth curves of mixed cultures of Pots (g) and L3.3 (v) strains. D) Fitness comparison of the two strains at different inoculation ratios. Single colony swim rings were easy to detect and isolate at 100:1 (Pots:L3.3) dilution (orange arrow). All growth curves are averages of 3 independent measurements.

A

| Primer ID # | Primer Name | Primer Sequence (5' to 3') | Annealing Temperature (°C) |
|---|---|---|---|
| 1 | E.Coli MotA/B_RA FW | GCGAAAGGATAATTCGTCG | 60 |
| 2 | E.Coli MotA/B_RA RV_oh:PotB | AACCGAGGTGACAGC | 60 |
| 3 | LA/pSHU_FW | GATGATGTCGTGGATTTAGCAACCC | 58 |
| 4 | RA/pSHU_RV | CTGTCACCTCGGTTCG | 58 |
| 5 | E.Coli MotA/B_LA FW_oh:PomA | CTAAATCCACGACATCATCCTTCCACTG | 60 |
| 6 | E.coli MotA_LA RV | TTATGACCTGGGAACAAAAC | 60 |
| 7 | E.Coli Xsome_1_FW | AGCCCGAAAGATGGCATTCA | 68 |
| 8 | E.Coli Xsome_2_FW | TCGCGCATATAGGGTTCGAC | 68 |
| 9 | E.Coli Xsome_3_RV | ACTTTCCCAGAATCCTGCCG | 68 |
| 10 | PomA_ RV | AATCAAACCTAATAGGGTTGCTAAATCCAC | 68 |
| 11 | E.Coli Xsome_4_ RV | ATGCTGCCATTCTCAACCGA | 68 |
| 12 | PotB G20V_g59t_RV | TCAAATCTGCGAATGTCACCATCCATAACGGGAGG | 55 |
| 13 | PotB G20V_g59t_FW | CCTCCCGTTATGGATGGTGACATTCGCAGATTTGA | 55 |

B

**Overlap Extension**

| PCR cycles | melting | annealing | extension |
|---|---|---|---|
| x1 | 98'C 90s | | |
| x30 | 98'C 20 s | X'C 30s | 72'C 1 min |
| x1 | | | 72'C 3 min |

**Assembly**

| PCR cycles | melting | annealing | extension |
|---|---|---|---|
| x1 | 98'C 2 min | | |
| x25 | 98'C 30 s | 57'C 1 min | 72'C 3.5 min |
| x1 | | | 72'C 5 min |

**Nested PCR**

| PCR cycles | melting | annealing | extension |
|---|---|---|---|
| x1 | 98'C 2 min | | |
| x25 | 98'C 30 s | 68'C 1 min | 72'C 3.5 min |
| x1 | | | 72'C 5 min |

**Colony PCR Screening**

| PCR cycles | melting | annealing | extension |
|---|---|---|---|
| x1 | 98'C 2 min | | |
| X34 | 98'C 30 s | 67'C 1 min | 72'C 1 min |
| x1 | | | 72'C 5 min |

**Colony PCR Sequencing Amplicon**

| PCR cycles | melting | annealing | extension |
|---|---|---|---|
| x1 | 98'C 2 min | | |
| X34 | 98'C 30 s | 61'C 1 min | 72'C 1 min |
| x1 | | | 72'C 5 min |

**Quikchange PCR protocol**

| PCR cycles | melting | annealing | extension |
|---|---|---|---|
| x1 | 98'C 30s | | |
| X34 | 98'C 30 s | 55'C 1 min | 72'C 5 min |
| x1 | | | 72'C 5 min |

**Supplementary Fig. 18. Primer list and PCR protocols.** A) List of primers used in this work. B) List of PCR protocols employed.

**Supplementary Table 1**

**SNP comparison of RP437 and MG1655 genomes.** The list indicates all the shared SNPs across RP437 descendants detected after WGS. The short reads from sequencing were aligned to the MG1655 reference *E. coli* genome (GenBank: U00096.2).

**Supplementary Table 2**

**RNAseq gene counts and DESEQ2 results**. Excel spreadsheet tabs contain: Gene counts and z-scores (list of all read counts per E. coli gene, for each experimental replicate and their associated z-scores); L183F heatmap scores (gene names and z-scores as displayed in Fig.2A heatmap); G20V heatmap scores (gene names and z-scores as displayed in Fig.2B heatmap); All pairwise analysis DESEQ2 results. DESEQ2_L3.2(preG20V) v Pots; L3.3(G20V) v L3.2(preG20V); L3.3(G20V) v Pots; L5.3(preL183F) v Pots; L6.4(L183F) v L5.3(preL183F); L6.4(L183F) v Pots; Genome Annotation table for Pots (GenBank: CP083410.1).

**Supplementary Table 3. Quantification of residual sodium in buffers and media.** Residual sodium levels (mM) in cell culture media components and motility buffers measured by Atomic Absorption Spectroscopy. N = 1 (tryptone, yeast extract, and agar), N=2 (Buffer only: 10 mM KPi + 85 mM KCl + 0.1 mM EDTA-2K , 10 mM KPi + 85 mM KCl; Values indicate mean + standard deviation, where there are two measurements.) *E. coli* RP437 Δ*motAB* Δ*cheY* Δ*pilA* *fliC*$^{st}$ (FliC-sticky) cells carrying pPots *pomApotB* (G20V) were grown at 30ºC in $Na^{+}$TB broth with 1 mM arabinose and 25 µg/mL chloramphenicol. The cells (1 mL) were harvested by centrifugation and washed with 1 mL $K^{+}$ motility buffer ($K^{+}$MB: 85 mM KCl, 10 mM KPi, 0.1 mM EDTA-2K, pH=7.0). This procedure was repeated one or three times then the rest of the cell suspension was centrifuged, and the sodium concentration of its supernatant was measured by AAS after calibration with standard sodium solutions of known concentrations. Cell culture media components (yeast extract, Bacto tryptone and Bacto agar) were diluted 100-1000x before AAS measurements. In tethered cell and swimming assays in this work three washes were performed

| Buffer Ingredients | |
|---|---|
| | **[Na+] (mM)** |
| **1% Bacto tryptone** | 12.8 |
| **0.5% yeast extract** | 1 |
| **0.3% Bacto agar** | 0.96 |
| **0.1M KCl** | 0.0157 |
| Buffers | |
| **$K^{+}$ Motility Buffer ($K^{+}$MB: 10 mM KPi, 85 mM KCl)** | |
| Buffer only | 0.020 |
| Cell harvest with 1x wash | 0.0325 ± 0.0025 |
| Cell harvest with 3x wash – used for assay data in Fig. 3 and Supplementary Fig. 10. | 0.023 ± 0.002 (N=4) |
| **10 mM KPi, 85 mM KCl, 0.1 mM EDTA-2K** | |
| Buffer only | 0.032 |
| Cell harvest with 1x wash | 0.044 ± 0.003 |
| Cell harvest with 3x wash – used for assay data in Supplementary Fig. 11 & 12 | 0.036 ± 0.003 (N=4) |
| Soft Agar Swim Plates | |
| **$K^{+}$LB** | ~15 |
| **$Na^{+}$LB** | ~100 |
| **$K^{+}$ Minimal Media** | ~1 |
| **$Na^{+}$ Minimal Media** | ~86 |

**Supplementary Video 1.** Motility of free-swimming *E. coli* Pots cells in $Na^+$MB. Original Resolution: 1224 x 1024 pixels. Field of view: 180 μm x 150 μm.

**Supplementary Video 2.** Motility of free-swimming *E. coli* Pots cells in $K^+$MB. Original Resolution: 1224 x 1024 pixels. Field of view: 180 μm x 150 μm.

**Supplementary Video 3.** Motility of free-swimming *E. coli* L3.3 cells (*potB* G20V) in $Na^+$MB. Original Resolution: 1224 x 1024 pixels. Field of view: 180 μm x 150 μm.

**Supplementary Video 4.** Motility of free-swimming *E. coli* L3.3 cells (*potB* G20V) in $K^+$MB. Original Resolution: 1224 x 1024 pixels. Field of view: 180 μm x 150 μm.

**Supplementary Video 5.** Motility of free-swimming *E. coli ΔmotAB* cells + pPots in $Na^+$MB. Original Resolution: 1224 x 1024 pixels. Field of view: 180 μm x 150 μm.

**Supplementary Video 6**. Motility of free-swimming *E. coli ΔmotAB* cells + pPots in $K^+$MB. Original Resolution: 1224 x 1024 pixels. Field of view: 180 μm x 150 μm.

**Supplementary Video 7.** Motility of free-swimming *E. coli ΔmotAB* cells + pPots *potB* G20V in $Na^+$MB. Original Resolution: 1224 x 1024 pixels. Field of view: 180 μm x 150 μm.

**Supplementary Video 8.** Motility of free-swimming *E. coli ΔmotAB* cells + pPots *potB* G20V in $K^+$MB. Original Resolution: 1224 x 1024 pixels. Field of view: 180 μm x 150 μm.
